# Cap-adjacent 2’-*O*-ribose methylation of RNA in *C. elegans* is required for postembryonic growth and germline development in the presence of the decapping exonuclease EOL-1

**DOI:** 10.1101/2025.03.10.638824

**Authors:** Eileen Clemens, Sarah Brivo, Mohammed Al-Khafaji, Peter Eijlers, Maheshika Kurukulasuriya, Irmgard U. Haussmann, David MacLeod, Marius Wenzel, Berndt Müller, Matthias Soller, Jonathan Pettitt

## Abstract

Cap-adjacent 2’-*O*-ribose methylation (cOMe) of the first two transcribed nucleotides of RNA polymerase II transcripts is a conserved feature in many eukaryotes. In mammals, these modifications are key to a transcript surveillance system that regulates the interferon response, but the broader functions of cOMe remain poorly understood. To understand the role of cOMe in *C. elegans*, we functionally characterised the methyltransferases (CMTR-1 and CMTR-2) responsible for installing these modifications. These enzymes have distinct expression patterns, protein interaction partners, and loss of function phenotypes. Loss of CMTR-1 causes dramatic reductions in cOMe, impaired growth and sterility. In contrast, animals lacking CMTR-2 are superficially wild-type, though CMTR-2 loss enhances the severity of the *cmtr-1* mutant phenotype. Depletion of CMTR-1 causes downregulation of transcripts associated with germline sex determination and upregulation of those involved in the intracellular pathogen response (IPR). We show that absence of the decapping exonuclease, EOL-1, an IPR component, completely suppresses the sterility and growth defects caused of loss of CMTR-1, suggesting that EOL-1 degrades cellular transcripts lacking cOMe. Our work shows the physiological relevance of cOMe in protecting transcripts from decapping exonucleases, raising the possibility that cOMe plays a role in RNA-mediated immune surveillance beyond the vertebrates.

**GRAPHICAL ABSTRACT:** 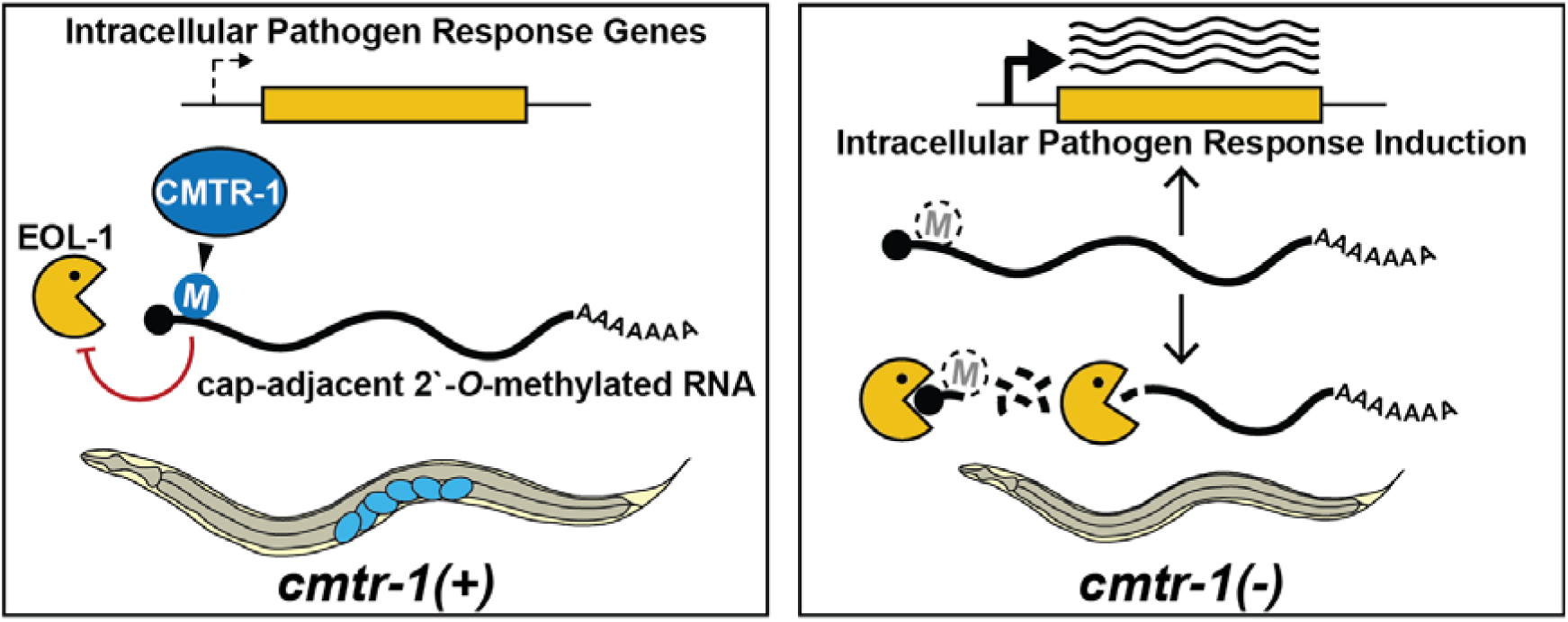

## INTRODUCTION

Cap-adjacent 2’-*O*-ribose methylation (cOMe) of RNA was first discovered in mammalian cells several decades ago (Adams and Cory, 1975; Anreiter et al., 2023; Furuichi et al., 1975; Galloway and Cowling, 2019). Subsequent work showed that these modifications are found throughout the Metazoa, as well as in the viruses and protists that infect these organisms. In animals, the first two transcribed nucleotides can harbour cOMe, with methylation of the first transcribed nucleotide being designated “cap1”, and methylation of both first and second nucleotides referred to as “cap2”. Although these modifications are ubiquitous, their physiological significance remains incompletely understood.

The first defined and the best understood role for cOMe is in preventing activation of an RNA-based immune response (Daffis et al., 2010; Li et al., 2013; Schuberth- Wagner et al., 2015; Züst et al., 2011): an antiviral mechanism consisting of sensor proteins that recognise transcripts lacking cOMe, which brings about the activation of the interferon response. This has created strong selective pressure for viruses to evolve the means to ensure their transcripts are modified by cOMe (Hyde and Diamond, 2015). However, this function does not explain the presence of cOMe in animals that lack the interferon pathway. It also doesn’t explain the full significance of cOMe in mammals: loss of either of the two highly conserved methyltransferases, CMTR1 and CMTR2, responsible for cOMe in mice results in embryonic lethality without inducing the interferon pathway (Dohnalkova et al., 2023); and there are multiple lines of evidence supporting diverse roles for cOMe, including the regulation of transcription, RNA splicing, and translation (Dönmez et al., 2004; Drazkowska et al., 2022; Haussmann et al., 2022; Kuge et al., 1998; Liang et al., 2022; Yu et al., 1998).

Studying cOMe in genetically tractable model organisms such as *Drosophila melanogaster* and *C. elegans* should allow us to understand the functional significance of cOMe from a whole animal perspective. Both organisms have orthologues of mammalian CMTR1 and CMTR2 (Dix et al., 2022; Haussmann et al., 2022). However, detailed functional genetic analysis in *Drosophila* has been hampered by the relatively subtle phenotypic consequences of loss of cOMe. *Drosophila CMTr1*/*2* double mutants are viable, though they show specific behavioural defects consistent with defects in the spatial regulation of translation in neurons (Haussmann et al., 2022).

Previous studies have shown that subcellular mislocalisation of *C. elegans* CMTR- 1can affect the expression of specific genes (Meisel et al., 2024). Here we present a comprehensive analysis of CMTR function, showing that loss of cOMe in *C. elegans* leads to growth retardation and pronounced defects in germline development, with *cmtr-1* loss-of-function mutants conferring a fully penetrant sterile phenotype. Loss of cOMe leads to activation of a set of genes that constitute the intracellular pathogen response (IPR), which are induced by viruses and intracellular pathogens (Lažetić et al., 2023), raising the possibility that as in mammals, there exists a surveillance mechanism to detect transcripts lacking these modifications. From a forward genetic screen for suppressors of *cmtr-1* loss-of-function we identified the decapping exonuclease EOL-1, showing that this enzyme is a major component of the cellular response to transcripts lacking cOMe. Our work suggests evolutionary continuity in the role of cOMe in RNA immune surveillance and provides physiological confirmation of the role of this modification in protecting transcripts from degradation by the DXO/Rai decapping exonuclease family.

## MATERIALS AND METHODS

### Nematode Strains

*C. elegans* strains were grown using standard culture conditions (Stiernagle, 2006) at 20°C unless otherwise indicated. N2 (Bristol) was used as the wild type. The following previously described strains were used in this study: HML1012 (Hills- Muckey et al., 2022); CGC45; CGC66. Strains generated in the current study are described in Supplementary Table 1.

The *cmtr-1* deletion allele, *syb3613*, was generated by Fujian Sunybiotech Co. LTD, and obtained as strain PHX3613. The *cmtr-2* deletion allele, *tm4453*, was obtained from the Mitani Lab as strain FX4453. PE1000 was generated from FX4453 by outcrossing six times against N2. Fluorescent-tag/degron knock-in strains were created as described previously (DeMott et al., 2021; Dickinson et al., 2015; Hills- Muckey et al., 2022). The CMTR-1(K244A) mutation was generated through oligonucleotide-templated repair of CRISPR/Cas9 induced double-strand breaks, using the previously described *dpy-10* based co-conversion strategy (Arribere et al., 2014). CRISPR alleles of *eol-1* were made using the same *dpy-10*-based approach, but the injections were performed using PE1176 animals and the resultant F1 progeny grown on 5-Ph-IAA containing plates, selecting for suppression of the sterile phenotype. Alleles were confirmed by Sanger sequencing (Eurofins Genomics).

Oligonucleotides used to verify the allelic status of genes, generate guide RNA constructs, and produce homologous repair templates are given in Supplementary Table 1.

### Transgenic rescuing assay

The wild-type *cmtr-1* cDNA was amplified from total RNA using primers cmcDNAFo and cmcDNARev (Supplementary Table 1). The plasmid backbone, which contains the *rps-0* promoter and *unc-54* 2’-O- UTR, was amplified using primers prpo-u54Fwd and prpo-u54Re (Supplementary Table 1), and the *cmtr-1* cDNA cloned between the *rps-0* promoter and *unc-54* 3’UTR using the NEBuilder HiFi DNA Assembly kit (NEB) to create plasmid pSB2. The plasmid was verified by Sanger Sequencing (Eurofins Genomics). PE1033 worms were co-injected with pSB2 and a plasmid containing *myo-2p::tdTomato* (Eijlers et al., 2024). Transgenic lines were established as previously described (Mello et al., 1991; Nance and Frøkjær-Jensen, 2019). The rescue assay was performed by picking non-tdTomato offspring as L1/L2 larvae to separate plates and scoring their ability to reach fertile adulthood.

### Fluorescent microscopy

Worms were mounted in 5 µl M9 supplemented with 10 mM sodium azide on 5% agar pads. Images were obtained using either a Zeiss Axioplan 2, equipped with a Hamamatsu Orca ER camera, or a Zeiss Imager M2 upright microscope, equipped with a Hamamatsu Flash 4 LT camera.

### Western blot

Western blots were performed as described previously (Fasimoye et al., 2022) using lysates from 300 – 500 adult/L4 larvae, or 5000 L1 larvae per well. To detect embryonic expression of CMTR-1/-2 in embryos, 40 µg of total protein from embryonic extracts was loaded per gel lane. For detecting larval expression, GFP-tagged proteins were detected with mouse anti-GFP antibodies (Roche 11814460001), mNeonGreen tagged proteins with rabbit anti-mNeonGreen antibodies (Proteintech, 29523-1-AP) and GAPDH using mouse anti-GAPDH antibodies (Invitrogen AM4300). Secondary antibodies used were anti-mouse or anti-rabbit HRP conjugated IgG antibody (Cell Signalling Technology 7074 and 7076). To detect proteins in embryonic extracts, GFP tag Polyclonal antibody (Proteintech, 50430-2-AP) (1:1000 dilution) was used together with the above anti-rabbit secondary (1:3000 dilution). Blots were also stained with amidoblack and quantified using either Fiji (Schindelin et al., 2012)or ImageQuant (Cytiva) software (Fasimoye et al., 2022).

### Preparation of *C. elegans* embryo extracts

*C. elegans* embryo extracts were prepared essentially as described previously (Eijlers et al., 2024)(see also Supplementary Methods). Extracts were treated with RNase as described previously (Eijlers et al., 2024).

### Cap-methylation status analysis of mRNA

Total RNA was extracted with Trizol (Sigma) according to the manufacturers description using 20 µg glycogen (Roche) for precipitation. poly(A) mRNA was prepared using the NEBNext® Poly(A) mRNA Magnetic Isolation Module (NEB) by double oligo dT selection according to the manufacturer’s instructions.

For the analysis of 5’ cap structures, 5 µg total RNA was used for decapping by yDcpS in 20 µl for 1 h at 37°C according to the manufacturer’s instruction (NEB). The RNA was extracted by phenol/CHCl_3_ and ethanol precipitated in the presence of glycogen. The RNA was then labeled in a total volume of 20 µl containing 2 µl capping buffer (NEB), 1 µl SAM (2 mM), 0.25 µl ^32^P-αGTP (3000 Ci/mmol, 6.6 µM; Hartmann Analytics), 0.5 µl RNase Protector (Roche) and 0.5 µl capping enzyme (NEB) by incubation for 1 h at 37°C. The volume was then increased to 50 µl with water and poly(A) RNA selected as described above. The RNA was then digested on the beads in 5 µl using 0.5 µl NEB buffer 3 and 0.5 µl RNase I for 2 h, and then 10 µl gel loading buffer was added (98% deionized formamide,10 mM EDTA, 0.025% xylene cyanol FF and 0.025% bromphenol blue); products were analysed on 22% denaturing polyacrylamide gels (National Diagnostics) and pre-run for 2 h. Gels were soaked in 20% PEG400, 10% acetic acid and 40% methanol for 10 min and then dried on a Whatman 3MM paper. Dried gels were then exposed to a storage phosphor screen (Bio-Rad) and scanned by a Molecular Imager FX in combination with QuantityOne software (Bio-Rad). Markers were prepared as described previously (Dix et al., 2022; Haussmann et al., 2022).

For the analysis of the first nucleotide in mRNA, 5 µg poly(A) mRNA was purified as described above and 5 µl incubated with terminator nuclease (Epicenter), according to the manufacturer’s instructions, to remove rRNA followed by phenol/CHCl_3_ and ethanol precipitation in the presence of glycogen (Roche). The mRNA was then decapped using RppH (NEB) and dephosphorylated by Antarctic phosphatase (NEB) in NEB buffer 2 supplemented with 0.1 mM ZnCl_2_ in 20 µl. Then the RNA was extracted by phenol/CHCl_3_ and precipitated in the presence of glycogen. The 5’-ends of dephosphorylated mRNAs were then labelled using 10 units of T4 PNK (NEB) and 0.25 µl ^32^P-γATP (6000 Ci/mmol, 12.5 µM; Hartmann Analytics). The labelled RNA was precipitated, and resuspended in 10 µl of 50 mM sodium acetate buffer (pH 5.5) and digested with nuclease P1 (SIGMA) for 2 h at 37°C. Two microliters of each sample was loaded on cellulose F TLC plates (20× 20 cm; Merck) and run in a solvent system of isobutyric acid:0.5 M NH_4_OH (5:3, v/v, solvent A), as first dimension, and isopropanol:HCl:water (70:15:15, v/v/v, solvent B) or sodium phosphate, 0.1 M (pH 6.8), ammonium sulfate, n-propanol (100:60:2, v/w/v, solvent C), as the second dimension. TLCs were repeated from biological replicates. The identity of the nucleotide spots was determined as described (Keith, 1995; Kruse et al., 2011). TLCs were exposed to a storage phosphorscreen (Bio-Rad) and scanned by a Molecular Imager FX in combination with QuantityOne software (Bio-Rad).

### Monitoring SL1 *trans*-splicing using quantitative PCR

RNA was isolated from embryonic extracts prepared from PE1176 and PE1220 embryos, treated for 4 h with either 50 µM 5-Ph-IAA-AM in DMSO, or control treated in DMSO by extraction with TriZol, and then purified using the PureLink RNA Mini kit (Life Technologies), with modifications for TRIzol treated samples and DNase treatment, as described by the manufacturer. Reverse transcription and analysis by qPCR were done as described (Eijlers et al, 2024, Philippe et al, 2017). Analyses were done as three technical replicates. Shown are ΔC_T_ values derived for each replicate (Schmittgen and Livak, 2008).

### *In vitro* spliced leader *trans*-splicing assays

For extract preparation, PE1176 and PE1220 animals were grown in liquid culture and embryos were isolated and treated with 50 µM 5-PH-IAA-AM or control-treated with DMSO as described (Eijlers et al., 2024), except that treatment was for 4h.

Extracts were then prepared as described (Eijlers et al., 2024) and stored at -80°C. The synthetic SL *trans*-splicing substrate, consisting of *rps-3* genomic DNA containing 404 bp of 5’ UTR sequence (including 382 bp of outron sequence upstream of the known SL1 *trans*-splice site) and extending 266 base pairs downstream from the ATG translation start site, was amplified from *C. elegans* genomic DNA using primers *rps-3*genFwd and *rps-3*genRev (Supplementary Table 1). The resulting amplicon was inserted into pBlueScript KS(-) cleaved with XbaI using NEB HIFI gene assembly to produce pBS-*rps-3*. The sequence was confirmed by Sanger sequencing (Eurofins Genomics). pBS-*rps-3* DNA linearised with EcoRI was transcribed with T7 RNA polymerase using the MEGAscript® kit (Invitrogen).

This produces a synthetic *rps-3* RNA with a 3’-end derived from pBlueScript KS(-) that serves as a primer binding site in PCR. RNA was extracted using phenol/chloroform, concentrated by ethanol precipitation, resuspended in RNase-free water and purified using MicroSpin G-25 columns (Cytiva). RNA concentration was determined by measuring absorption at 260 nm assuming 1 AU = 40 µg/ml.

*In vitro* SL1 *trans*-splicing reactions were done essentially as described (Lasda et al., 2011). 15 μl reactions were done in 10 mM Tris-HCl (pH 8.0), 60 mM KCl, 4 mM MgCl_2_, 2 mM ATP, 20 mM creatine phosphate, 50 μg/ml creatine kinase, 2 mM DTT, 3 % PEG 8000, 0.25 mM EDTA, 5% glycerol, 0.5 mM PMSF, 2 U/μl RNaseOUT, 8 µg/μl embryonic extract and 2.5 ng/μl synthetic *rps-3* mRNA. Control reactions were performed without addition of ATP, creatine phosphate and creatine kinase.

Incubation was at 15°C for 2 h.

For analysis by one-step RT-PCR, reactions were diluted 1:80 in RNase-free MilliQ water and aliquots were then used for one-step RT-PCR using the Luna® Universal One-Step RT-qPCR kit (NEB) according to the manufacturer’s instructions; primers are given in Supplementary Table 1. Reactions were done with 1 µl or 2 µl diluted reactions and analysed by agarose gel electrophoresis, or quantified using a LightCycler 480 (Roche), with software release 1.5.1.62 Sp3. qPCR data was analysed using the ΔC_T_ method (Schmittgen and Livak, 2008). Products P1, P2 and P3 were identified by cloning into pGEM-T Easy (Promega) followed by Sanger sequencing (Eurofins Genomics).

### Protein Immunoprecipitation

Immunoprecipitations of GFP-tagged proteins were performed in triplicates or quadruplicates using anti-GFP nanobody coupled agarose beads and control agarose beads (GFP-Trap and control agarose beads, Chromotek GmbH), essentially as described previously (Eijlers et al., 2024).

### Protein analysis by LC-MS/MS and differential protein expression analysis

Proteomic analysis by mass spectrometry was done at the Aberdeen Proteomics unit as previously described (Eijlers et al., 2024; Fasimoye et al., 2022). MaxQuant version 1.6.5.0 (Cox and Mann, 2008) was used to process the raw data files with the *C. elegans* reference proteome UP000001940 downloaded on the 24^th^ of August 2021 as described (Eijlers et al., 2024), except that the feature “match between runs” was applied. The MaxQuant protein group files were further analysed using Perseus version 1.6.5.0 as described (Eijlers et al., 2024). Graphs were drawn in GraphPad Prism.

### RNA extraction and mRNA-Seq

Prior to RNA extraction, the worms were treated with 5-Ph-IAA when they reached the L3 larval stage. Therefore, synchronised PE1176 L1s were grown in liquid culture for 22.5 h until they started moulting into larval stage 3. They were then divided into two and plated on control plates or 5-Ph-IAA plates, respectively, and cultured for 16 h, at which point the animals had started moulting into the L4 larval stage. Worms were washed off NGM plates using M9 and washed 3 times by centrifugation at 749g for 2 min at 4°C. Subsequently, 1 ml of Trizol (TRI Reagent, T9424, Sigma Aldrich) was added per 50 µl of worm pellet and samples frozen in liquid nitrogen. Samples were thawed at 37°C and re-frozen four times. They were then vortexed for 30 s, left to rest for 30 s and the vortex/rest process repeated for a further three times. Phase separation by addition of chloroform and recovery of RNA was done as described by the manufacturer. The aqueous fraction was transferred to a low DNA-binding-tube and diluted 1:1 with 70% ethanol. RNA was purified using PureLink RNA Mini Kit (Thermo Fisher Scientific), using the on-column DNase-treatment option. Three libraries per treatment group were sequenced on an Illumina NovaSeq instrument, generating 2x150bp paired-end reads (Novogene UK Ltd).

### Differential gene expression analysis

The quality of the raw reads was inspected in FASTQC 0.11.9 (Andrews, 2010) and MULTIQC 1.12 (Ewels et al., 2016), and bases with a Phred score below 20 were trimmed using TRIMGALORE 0.6.6 (Krueger, 2015). The trimmed reads were then aligned to the *C. elegans* WB235 reference genome using HISAT2 2.2.0 (Kim et al., 2015). Alignments were processed using SAMTOOLS 1.14 (Li et al., 2009). Gene-level read counts were obtained using FEATURECOUNTS 2.0.2 (Liao et al., 2014), quantifying against exon annotations and assigning fractional counts to all alignment locations of multi-mapping reads.

The read counts were analysed in DESeq2 1.42. (Love et al., 2014), identifying differentially expressed genes with an FDR (Benjamini and Hochberg, 1995) significance threshold of 0.05, and shrinking fold changes at low-count genes using the *apeglm* method (Zhu et al., 2019). Significantly differentially expressed genes (DEGs) were examined for overrepresentation GeneOntology terms (biological process ontology) using CLUSTERPROFILER 4.10.0 (Yu et al., 2012) with the *org.Ce.eg.db* 3.18.0 database (June 2024). The GO results were simplified by removing terms at graph levels 1-3, clustering GO terms semantically with the Wang method at a similarity cutoff of 0.7 (Yu et al., 2010), and retaining for each cluster the single GO term with the smallest *P*-value.

### Data analysis using RStudio

RStudio was used for data analysis and graph generation (http://www.rstudio.com/). The following R packages were used: ggplot2 (Wickham, 2016), dplyr (https://dplyr.tidyverse.org/), carData/car (10.32614/CRAN.package.car), multcomp (10.32614/CRAN.package.multcomp), ggpubr (https://rpkgs.datanovia.com/ggpubr/), rstatix (https://rpkgs.datanovia.com/rstatix/), tibble (https://tibble.tidyverse.org/), RcolorBrewer (10.32614/CRAN.package.RColorBrewer).

### Screen for suppressors of *cmtr-1* loss-of-function

PE1176 worms were subjected to a standard ethyl methylsulfonate (EMS) mutagenesis. Thirty F1 cultures were established by subjecting the mutagenised parental animals to alkaline hypochlorite treatment and plating 500 F1 animals per 9 cm plate. The mutagenised animals and their F1 progeny were propagated on standard NGM agar plates. The F2 embryos from alkaline hypochlorite treated F1s were plated onto 9 cm NGM plates supplemented with 1 µM 5-Ph-IAA. Suppressor mutants, identified on the basis of presence of eggs in the uterus of adult F2s, were picked to establish lines, taking care to only maintain one line for each F1 lineage.

### Genetic analysis of suppressor mutants

We reasoned that our screen would generate two classes of fertile mutants: those that are *bona fide* extragenic suppressors of mNG^AID::CMTR-1 knockdown; and loss-of-function mutants in the *TIR1* transgene, which would lead to 5-Ph-IAA-independent expression of mNG^AID::CMTR-1. We thus screened each suppressor line for the presence of nuclear mNeonGreen fluorescence when grown on media containing 5-Ph-IAA. Mutant lines that showed mNeonGreen fluorescence in the presence 5-Ph-IAA were assumed to be *TIR1* mutants; the *TIR1* coding region for a subset of these was sequenced to confirm this assumption. Amplicons generated from single-worm PCRs using primer pairs TIR1Fwd+TIR1Rev1 and TIR1fo1+TIR1re2 were sequenced using the same oligos to prime Sanger sequencing reactions (Eurofins Genomics).

Mutants that lacked nuclear mNeonGreen fluorescence in the presence of 5-Ph-IAA were assumed to be genuine epistatic suppressors and were backcrossed with non-mutagenised PE1176. The resultant strains were crossed into the *cmtr-1(syb3613)* background to determine the ability of the suppressor mutations to suppress the sterile phenotype of the *cmtr-1* null allele.

### Sib-Selection and DNA Preparation

Two suppressor strains, PE1237 (*fe152*) and PE1245 (*fe154*), were subject to a sibling subtraction/whole genome sequencing method adapted from a previously reported method (Joseph et al., 2017). Male PE1053 worms were crossed with hermaphrodites from PE1237/1245. F1 progeny carrying the *feEx342* transgenic array were identified on the basis of pharyngeal tdTomato expression, picked to separate plates, and allowed to segregate F2 progeny. Two hundred *feEx342* transgenic F2s were picked singly to separate 35 mm plates and allowed to propagate. Homozygous suppressor lines (Sup) were identified on the basis of the presence of fertile non-tdTomato F3/F4 worms, and the absence of non-tdTomato “scrawny” F3/F4 worms, the latter being observed on plates whose frequency suggested that these were broods of suppressor heterozygotes. Lines that failed to inherit suppressor mutations (Nonsup) were identified on the basis of the complete absence of fertile non-tdTomato F3/F4 worms. Contaminated plates were discarded. Worms were washed off plates, and Sup/Nonsup cultures pooled as outlined previously (Doitsidou et al., 2010). The number of 35 mm F2 plates used to generate the Sup and Nonsup pools were as follows: *fe152* Sup = 16; *fe152* Nonsup = 19; *fe154* Sup = 19; *fe154* Nonsup = 19. Genomic DNA was prepared using the Qiagen Puregene Tissue Kit (Qiagen, 158063) as described previously (Doitsidou et al., 2010). One library per pool was sequenced on an Illumina NovaSeq X Plus instrument, generating 2x150bp paired-end reads (Novogene UK Ltd).

### Variant Detection

Quality control of the raw reads was carried out using FASTQC 0.11.9 (Andrews, 2010) and MULTIQC 1.12 (Ewels et al., 2016). Adaptor removal and trimming of bases with a Phred score below 20 was performed using TRIMGALORE 0.6.6 (Krueger, 2015). The trimmed reads were then aligned to the *C. elegans* WB235 reference genome using BWA MEM 0.17.17 (Li and Durbin, 2010). Variant detection was done for all libraries together in a single run of FREEBAYES 1.3.6 (Garrison and Marth, 2012), and the resultant VCF files were filtered and manipulated using BCFTOOLS 1.14 (Danecek et al., 2021). We considered biallelic SNPs with experiment-wide genotyping quality of at least 20 and at least 10 reads in each library, and retained SNPs fixed for the alternative allele (>90% of read depth) in Sup libraries and fixed for the reference allele (>90% of read depth) in Nonsup libraries. SNPEFF 4.3t (Cingolani et al., 2012) was then used to functionally annotate the candidate SNPs identified by this filtering strategy.

## RESULTS

### CMTR-1 is required for oocyte cell fate

To determine the role of cOMe in *C. elegans,* we studied the phenotypes of animals lacking one or both CMTR proteins. We obtained a deletion allele, *syb3613*, of *cmtr- 1* (SunyBiotech), which removes the methyltransferase domain, and is thus a predicted null allele. We also studied a deletion allele of *cmtr-2*, *tm4453*, that deletes key residues in the methyltransferase domain (Dix et al., 2022). While *cmtr- 1(syb3613)* homozygotes were able to reach adulthood, they were 100% sterile (Figure 1A,B) and were thinner than similarly staged *syb3613* heterozygotes.

**Figure 1.**
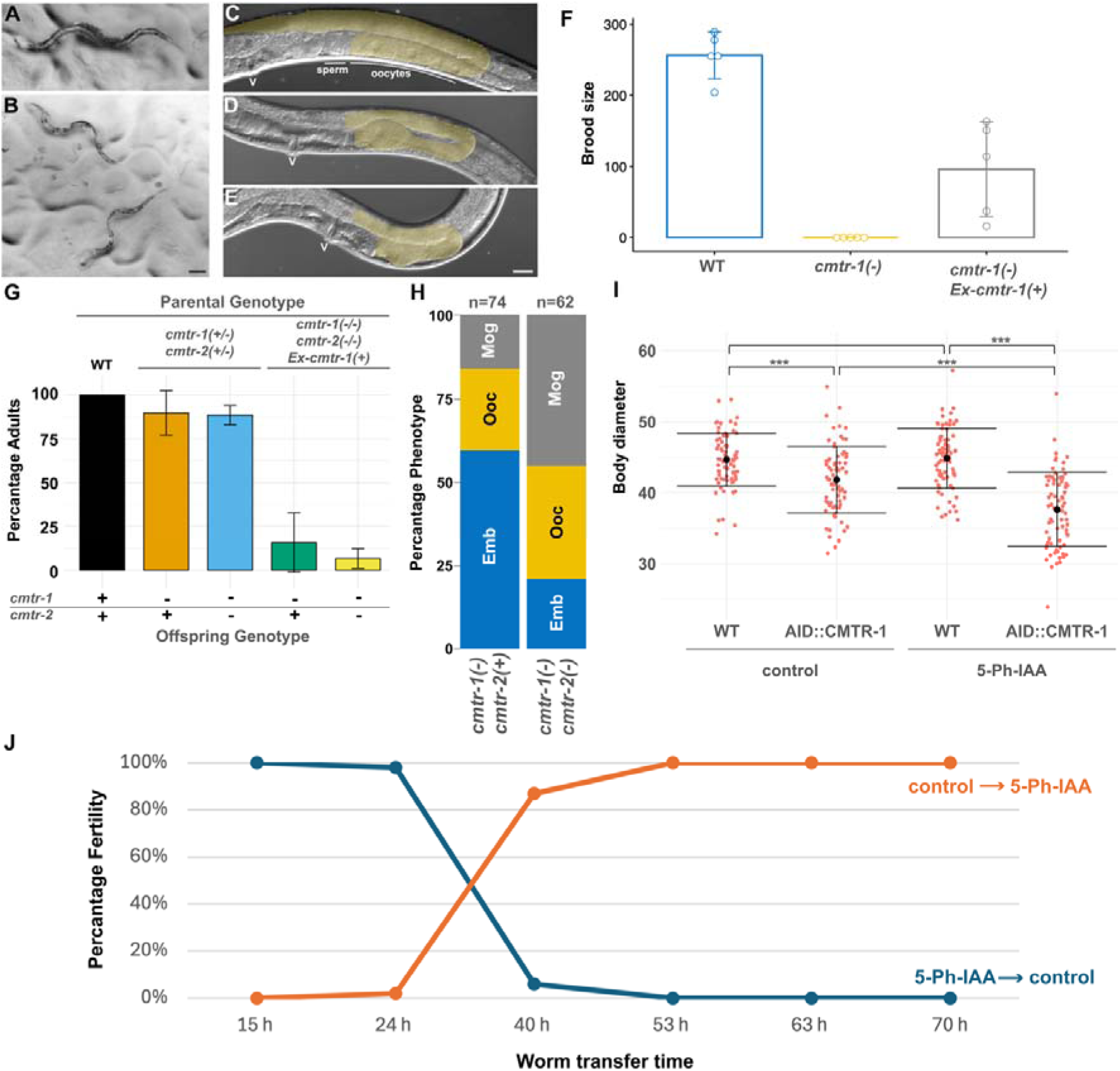
Loss of *cmtr-1* function results in defects in germline development and body size. Representative *cmtr-1(syb3613)/+* heterozygote (A), compared to sibling *cmtr-1(syb3613)* homozygotes (B). Wild type young adult hermaphrodite (C), showing posterior gonad arm (gold overlay), compared to the same regions of *cmtr- 1(syb3613)* homozygotes, one of which shows a single defective oocyte (D), and the other has a fully masculinised germline that has only produced sperm (E). Position of the vulva is indicated by “v”. Scale bars represent 100 µm (A,B) and 20 µm (C,D,E). (F) Transgenic expression of *cmtr-1* cDNA under the control of the *rps-0* promoter (*Ex-cmtr-1(+)*) rescues the fertility defects of *cmtr-1(syb3613)* mutants (*cmtr-1(-)*). (G) Comparison of the number of worms reaching adulthood 72 hours after hatching (n > 61 for each experiment). Mutant offspring (*cmtr-1(-)*; *cmtr-2(+)* or *cmtr-1(-)*; *cmtr-2(+)*) of transgenic parents (transgene is indicated ‘*Ex-cmtr-1(+)*)are slower-growing compared to genetically identical offspring of heterozygous parents. (H) Loss of *cmtr-2* increases the severity of the *cmtr-1* loss-of-function germline defects. Histograms show the distribution of the three germline defect classifications caused by loss of *cmtr-1* function: production of a few arrested embryos (Emb, blue); abortive oocyte production (Ooc, gold); masculinised germline (Mog, grey). (I) Worm diameter, measured at the mid-body, for worms homozygous for the mNG^AID::*cmtr-1* allele subject to control and 5-Ph-IAA treatment. Two-way ANOVA and a Tukey post hoc test were used to determine p-values; * < 0.05, ** < 0.01, *** < 0.001, n = 75 animals per group. (J) “Auxin-shift” experiment to determine when CMTR-1 function is required for fertility. Starved L1 larvae (n=50) were placed on either 5-Ph-IAA or control plates for the indicated time period, before being transferred to the alternative plate type (i.e. those on 5-Ph-IAA plates were moved to control plates, and vice versa). The worms were then left on the second plate, allowed to reach adulthood, and scored for fertility.

We found that the sterility was due to defects in the formation of oocytes, which ranged from a completely masculinised germline that produced only sperm, through to animals that were able to produce only a few embryos that failed to develop (Figure 1D,E,H). In all cases, *cmtr-1(syb3613)* homozygotes showed much smaller germlines than wild-type worms, something also observed in mutants that cause masculinised hermaphrodite germlines (Kimble and Crittenden, 2007).

We confirmed that the defects observed were due to loss of *cmtr-1* function by demonstrating that an extrachromosomal transgene consisting of the *cmtr-1* cDNA expressed under the control of the *rps-0* promoter rescued the germline sterility defects (Figure 1F,G). As part of this analysis we found that offspring that failed to inherit the extrachromosomal transgene generally showed a more severe phenotype than the offspring of heterozygous mothers, displaying more severe growth defects (Figure 1G). Since extrachromosomal transgenes are generally silenced in the germline (Kelly et al., 1997), this likely indicates that there is a significant *cmtr-1* maternal contribution, which is absent from the non-transgenic offspring of transgenic mothers. The more severe phenotypes of these animals indicates that the loss of *cmtr-1* function impacts other tissues beyond the germline.

In contrast to loss of *cmtr-1*, *cmtr-2* loss-of-function homozygotes are superficially wild type (Dix et al., 2022). To investigate whether loss of *cmtr-2* function influenced the *cmtr-1* loss-of-function phenotype, we examined the phenotype of *cmtr-1*; *cmtr-2* double mutants. This analysis showed that loss of *cmtr-2* enhanced the phenotype of *cmtr-1* single mutants (Figure 1H). Thus, CMTR-2 contributes to the role played by CMTR-1.

The failure of oocyte differentiation in *cmtr-1* mutant homozygotes suggested that CMTR-1 function is required for some component(s) of the sperm-oocyte switch. To determine when CMTR-1 protein is required during larval development for oocyte cell fate, we generated an auxin-inducible degeneration (AID) allele of *cmtr-1*, which expresses endogenous CMTR-1 tagged with the auxin-inducible degron (AID) peptide and monomeric NeonGreen (mNG^AID::CMTr1). This strain (PE1176) also expresses the *A. thaliana* TIR1(F79G) transgene necessary for auxin-dependent protein depletion (Hills-Muckey et al., 2022). Growing this strain in the presence of 5- Ph-IAA showed rapid depletion of mNG^AID::CMTr1 (Supplementary Figure 1), and such worms were 100% sterile and showed reduced body size compared to untreated controls (Figure I).

Using this strain, we performed a series of experiments designed to deplete mNG^AID::CMTR-1 at specific times during postembryonic development. We initiated synchronous cultures of 50 starved, first-stage larvae (L1s) on control plates and transferred them to 5-Ph-IAA plates at defined times (Figure 1J). We also performed the inverse experiment, culturing L1s on 5-Ph-IAA plates and transferring them to control plates at the same time points. This “auxin-shift” experiment (named by analogy to temperature-shift experiments performed with temperature-sensitive mutants) revealed that 5-Ph-IAA depletion of mNG^AID::CMTR-1 resulted in sterility only during a critical time period between approximately 24 and 40 hours following initiation of starved L1 larval development. The critical period during which CMTR-1, and thus cOMe, is required corresponds roughly to the L3 and L4 larval stages.

Given that the sperm-oocyte switch occurs during the L4 stage (Kimble and Crittenden, 2007), the time-window during which CMTR-1 is required suggest that this is when transcripts involved in germline sex determination need to be modified by cOMe for normal germline cell fate.

### The *C. elegans* CMTR proteins show distinct expression patterns and protein- protein interactions

Studies in cultured mammalian cells and *Drosophila* indicated that the two CMTR proteins have conserved, distinct subcellular distributions: in both cases, CMTR1 is primarily nuclear, whereas CMTR2 is both nuclear and cytoplasmic (Haline-Vaz et al., 2008; Haussmann et al., 2022; Werner et al., 2011). Using strains that express GFP- tagged alleles of the endogenous proteins, we found that *C. elegans* CMTR proteins are nuclear throughout postembryonic development (Figure 2A-D), being expressed in most, if not all cells, though CMTR-2 does not appear to be as strongly expressed in the nucleus as CMTR-1. While the overall levels of the two proteins are similar in embryos, as judged by Western blot (Figure 2H), the intracellular localisation of the two proteins is strikingly distinct: CMTR-1 is predominantly nuclear (Figure 2E), in contrast to CMTR-2, which is strongly cytoplasmic (Figure 2F, compare to Figure 2G).

**Figure 2.**
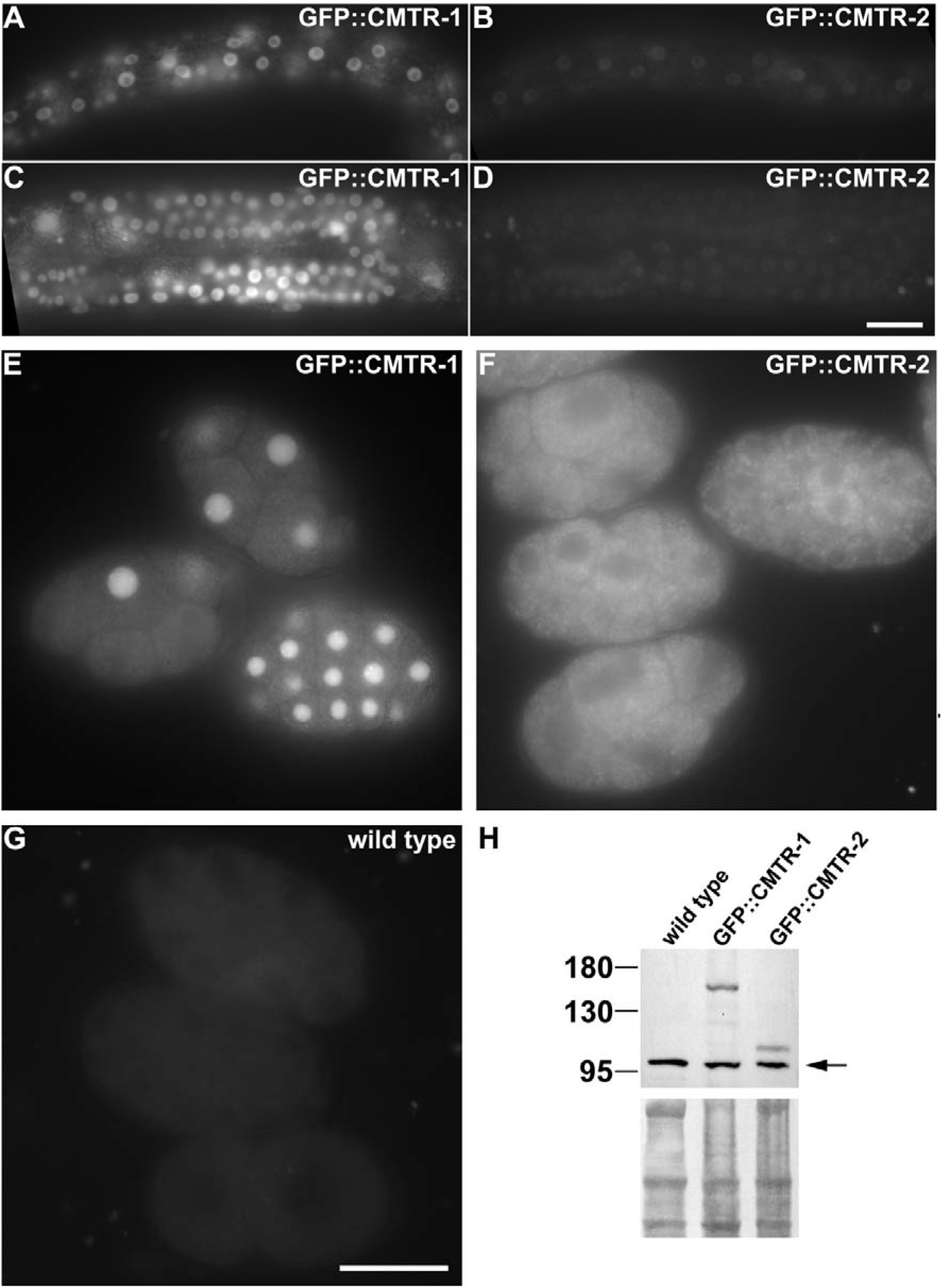
CMTR-1 is constitutively nuclear but CMTR-2 subcellular localisation is developmentally regulated. Representative images showing the respective expression of GFP tagged CMTR-1 (A, C, E) or CMTR-2 (B, D, F) in L2 larval stage epidermis (A, B), L4 germlines (C, D) and early embryos (E, F). Wild type embryos (G) show the level of embryonic background autofluorescence. Images in A – D and E - G were taken at the same camera exposure settings. Scale bars indicate 20 µm. (H) Western blot of GFP-tagged CMTR-1 and CMTR-2 in embryonic extracts probed with anti-GFP polyclonal antibody. Specific signals are absent from wild type embryos. Arrow indicates non-specific primary anti-GFP antibody immunoreactivity. Amido black stained immunoblot indicates loading of protein in each lane.

To investigate the functional significance of the distinct localisation of the proteins during embryogenesis further, we performed immunoprecipitation analyses from embryonic lysates to assess their respective protein-protein interactions. This analysis revealed that the two proteins have non-overlapping protein partners: CMTR-1 showed a rich set of protein-protein interactions (Figure 3A-C); while CMTR-2 displayed a paucity of interactors (Figure 3D, E); none of them matching those we found for CMTR-1. The lack of overlap in protein interaction partners is consistent with the low levels of CMTR-2 in the nucleus (Figure 2E,F) in embryos, but this analysis fails to shed light on the functional significance of the cytoplasmic enrichment of CMTR-2.

**Figure 3.**
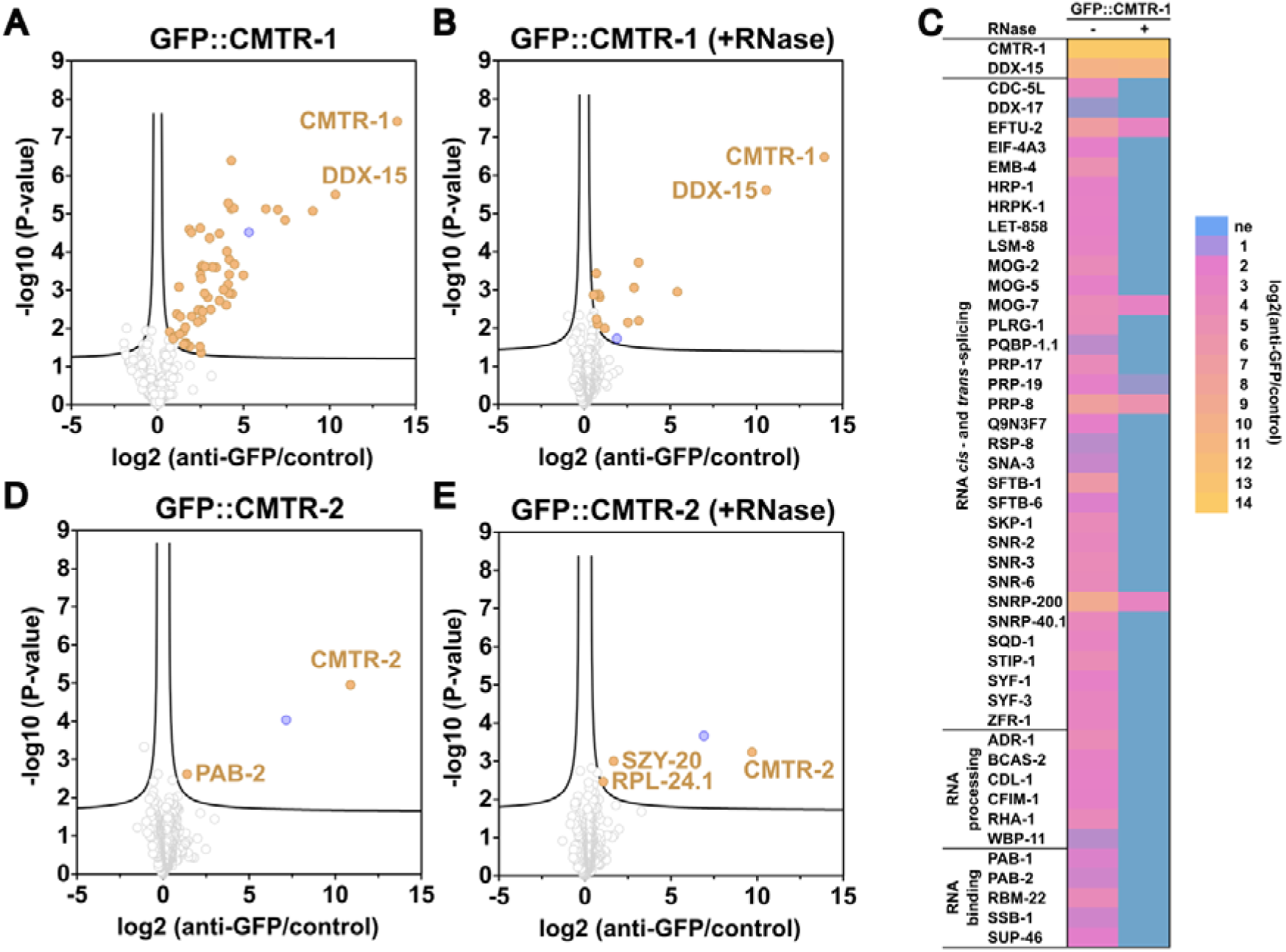
CMTR-1 and CMTR-2 show distinct protein-protein interaction partners. Proteins enriched by co-immunoprecipitation with GFP::CMTR-1 (A, B, C) and GFP::CMTR-2 (D, E) from PE1001 and PE991 embryonic extracts treated without (A, D) and with RNase (B, E). Immunoprecipitations were performed in triplicate or quadruplicate using anti-GFP nanobodies coupled to agarose beads, or control agarose beads. Immunoprecipitated proteins were identified by label-free quantification as described (Eijlers et al 2024). Graphs show enrichment in anti-GFP nanobody immunoprecipitations (anti-GFP) compared to bead only controls (control) and the false discovery rate (y-axis). Proteins that were significantly enriched are shown in gold. The significance cutoff curve is drawn in black (false discovery rate <=0.05 and S0 of 0.1). TBH-1 (blue), a dopamine β-hydroxylase, is found in all immunoprecipitations but our previous work suggests that it non-specifically associates with GFP-tagged proteins (Eijlers et al., 2024; Fasimoye et al., 2022). (**C**) Heatmap summarising the enrichment of spliceosome components, RNA processing factors and RNA binding proteins in immunoprecipitation of GFP::CMTR-1. Proteins were grouped using the gene ontology annotation retrieved from UniprotKB (Ashburner et al., 2000; Gene Ontology Consortium et al., 2023; The UniProt Consortium et al., 2023). ne: not enriched. For complete lists of identified proteins see Supplementary Table 2.

*C. elegans* CMTR-1 showed broadly the same protein-protein interactions with a suite of pre-mRNA processing factors that were observed for mammalian cells (Simabuco et al., 2019), with components of the spliceosome, heterogeneous nuclear ribonucleo-proteins and ribosomal proteins being prominent interaction partners (Figure 3A-C). We also saw an interaction with SNA-3, a component of the spliced leader *trans*-splicing machinery (Fasimoye et al., 2022) that has previously been shown to associate with nascent RNA polymerase II transcripts (Eijlers et al., 2024). Many of the interactions were abolished by RNase treatment, consistent with CMTR-1 interacting with them by virtue of binding to the same transcript. However, there are notable interactions that were preserved in RNase treated samples, included the previously well-defined CMTr1 interacting protein, DHX15 (DDX-15 in *C. elegans*)(Inesta-Vaquera et al., 2018; Toczydlowska-Socha et al., 2018), and some core components of the spliceosomal catalytic complex.

### Loss of cOMe is the cause of the *cmtr-1* loss-of-function phenotype

The phenotypic impact of loss of *cmtr-1* compared to *cmtr-2* suggested that CMTR-1 function might be more important for cOMe modification of transcripts than CMTR-2. This was supported by our previous analysis, which showed that loss of *cmtr-2* did not detectably affect cOMe levels (Dix et al., 2022). To investigate the dependence of cOMe on CMTR-1, we assayed RNA derived from animals in which mNG^AID::CMTR-1 was knocked down on its own, and in combination with the *cmtr- 2(tm4453)* loss-of-function mutation. Using a sensitive assay based on recapping mRNA with ^32^P-alpha-GTP, followed by RNase I digestion to detect cOMe in small amounts of mRNA (Anreiter et al., 2020; Dix et al., 2022; Haussmann et al., 2022), we found that cOMe modified RNA (cap1) was almost completely abolished (Figure 4A). Note that using this assay, we were not able to detect cap2 modifications, consistent with a previous study that this cap2 is largely absent from *C. elegans* (Despic and Jaffrey, 2023). This assay also revealed that CMTR-2 makes a negligible impact on cap1 levels, as we previously demonstrated (Dix et al., 2022).

**Figure 4.**
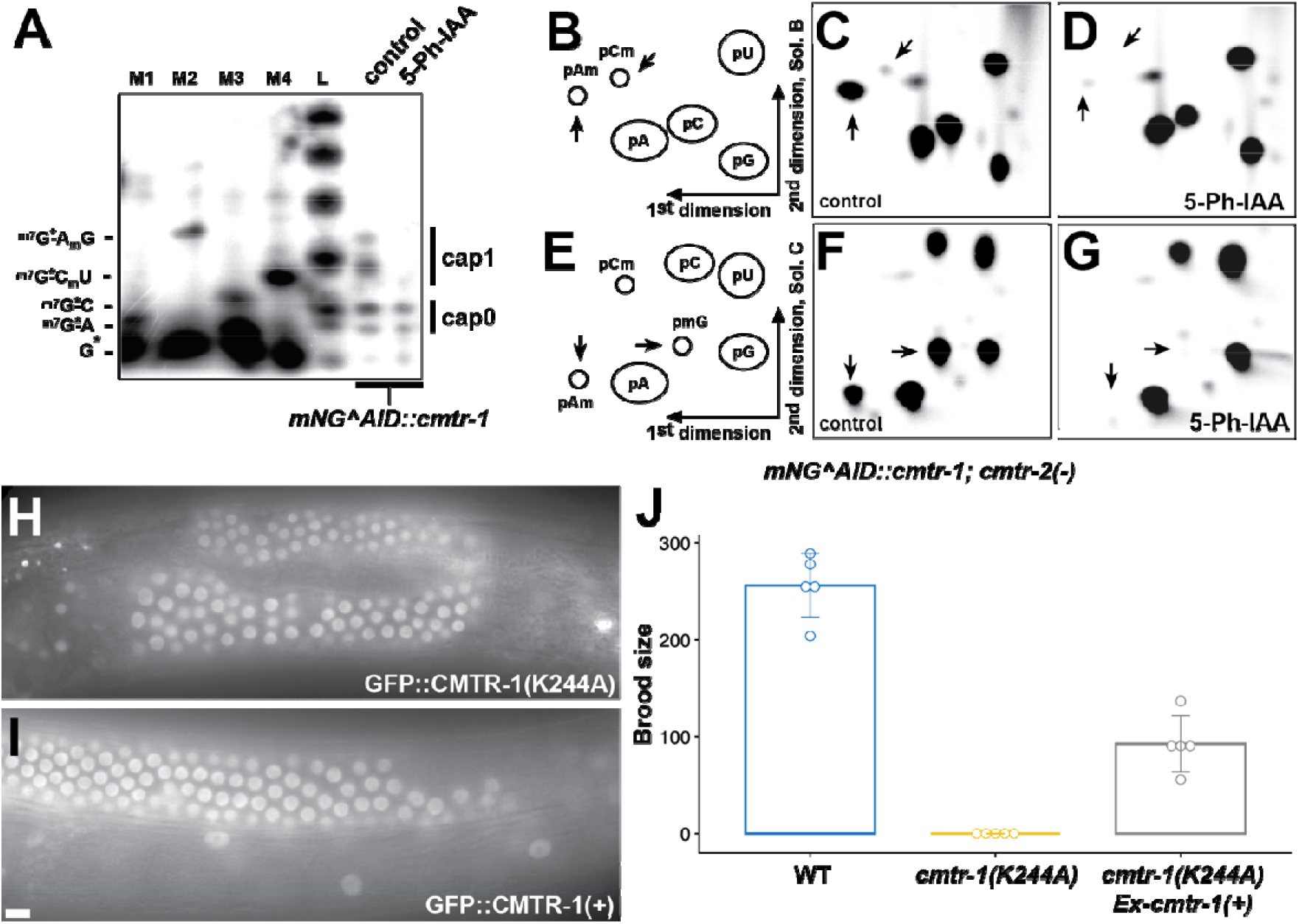
CMTR-1 is required for cap-adjacent 2’-O-ribose methylation in *C. elegans*. Analysis of cap-adjacent 2’-*O*-ribose RNA methylation with (control) and without (5-Ph-IAA) mNG^AID::CMTR-1. (A) Recapping assay to detect mRNA using ^32^P-αGTP. 5′ cap structures were separated on a 22% denaturing polyacrylamide gels after digestion with RNase I (lanes 6-7, right). Markers (M1-4) are RNase I digested ^32^P-αGTP capped oligonucleotides. 2’-*O*-ribose methylation (N_m_) was added using vaccinia CMTr. Sequences of markers are shown on the left. (B - G) Two dimensional thin-layer chromatograms (2D-TLCs) showing modifications of the first cap-adjacent nucleotides prepared from total RNA isolated from worms homozygous for the mNG^AID::*cmtr-1* and *cmtr-2(-)* alleles, grown on control (C, F) or 5-Ph-IAA plates (D, G). (B, E) Schematic diagrams of a 2D-TLC, depicting the positions of unmodified and 2’-*O*-ribose methylated nucleotides. One set of chromatograms (C, D) was run in solvent B for the second dimension, another set (F, G) was run in solvent C. 2’-O-ribose methylated nucleotides are indicated by arrows. (H) Representative germline GFP fluorescence of an animal expressing GFP-tagged CMTR-1(K244A), compared to GFP-tagged wild-type CMTR-1 protein. (J) Brood counts of wild-type (data from Figure 1F) and animals homozygous for the K244A mutation, in the absence and presence of the wild-type *cmtr-1* cDNA expressed under the control of the *rps-0* promoter (*Ex-cmtr-1*). Counts were performed on five individual animals per genotype.

Thus, CMTR-1 is the important determinant of cOMe in *C. elegans*, consistent with its strong loss-of-function phenotype.

We then looked specifically at cap1 in poly(A) mRNA derived from animals lacking functional CMTR-2 and depleted of mNG^AID::CMTR-1 by treatment with 5-Ph-IAA to find that cOMe is essentially absent (Figure 4B-G). Of note, the first transcribed nucleotide for the majority of *C. elegans* mRNAs will be the guanosine originating from the *trans*-spliced leaders. As shown by separation of the second dimension in solvent C, the spliced leader is also 2’-*O*-ribose methylated by CMTR-1 (Figure 4E- G), as expected for an RNA polymerase II transcript.

Previous studies in mammals have shown that around 50% of CMTR1 is in a complex with DHX15, and that this interaction represses its methyltransferase activity (Inesta-Vaquera et al., 2018; Toczydlowska-Socha et al., 2018). CMTR1 also influences RNA polymerase II recruitment to transcription start sites and the transcription of histone and ribosomal protein genes (Liang et al., 2022). We thus wanted to confirm that the *cmtr-1* loss-of-function phenotype was due solely to loss of cOMe; in other words that it was dependent on CMTR-1 methyltransferase activity.

To address this, we engineered a missense mutation, K244A, into the endogenous *cmtr-1* gene. The orthologous mutation in human CMTR1 (K239A) has previously been show to abolish methyltransferase function (Bélanger et al., 2010; Feder et al., 2003), and the lack of function of this mutation in *C. elegans* has been assessed in transgene rescue experiments (Meisel et al., 2024). We carried out site-directed mutagenesis of the *cmtr-1* allele that expresses CMTR-1 with a N-terminal GFP tag (Figure 2), allowing us to monitor the impact of the K244A mutation on the expression and localisation of CMTR-1 (Figure 4H,I). All worms homozygous for the K244A (*fe175*) allele showed the same recessive sterile and growth defective phenotypes displayed by the *syb3613* mutation, and these phenotypes were complemented by the *rps-0::cmtr-1(+)* transgene (Figure 4J). Since these worms specifically lack the methyltransferase activity of CMTR-1, this indicates that loss of cOMe is the cause of the germline phenotypes.

### Loss of CMTR-1 does not affect spliced leader *trans*-splicing

Approximately 85% of *C. elegans* genes produce transcripts that are *trans*-spliced to the SL1 or SL2 spliced leaders (Tourasse et al., 2017), and both spliced leaders also possess cOMe modifications (Figure 4E-G). In addition, we observed an interaction between CMTR-1 and SNA-3, an essential component of the SL1 *trans*-splicing machinery (Fasimoye et al, 2022, Eijlers et al, 2024) (Figure 3C). We therefore wanted to investigate whether cOMe had any functional impact on *trans*-splicing. We depleted mNG^AID::CMTR-1 in embryos by treating them with 5-Ph-IAA-AM, which is able to penetrate the egg-shell (Negishi et al., 2022) and prepared *in vitro* splicing extracts as described previously (Lasda et al., 2011). These were compared to extracts prepared from untreated embryos, and to PE1220 embryos depleted for SNA-3::AID^mNG, which, based on SNA-3’s essential role, is predicted to impair spliced leader *trans*-splicing (Fasimoye et al., 2022). 5-Ph-IAA-AM treatment reduced mNG^AID::CMTR-1 and SNA-3::mNG^AID steady-state levels to undetectable levels and by 80%, respectively (Figure 5A).

**Figure 5.**
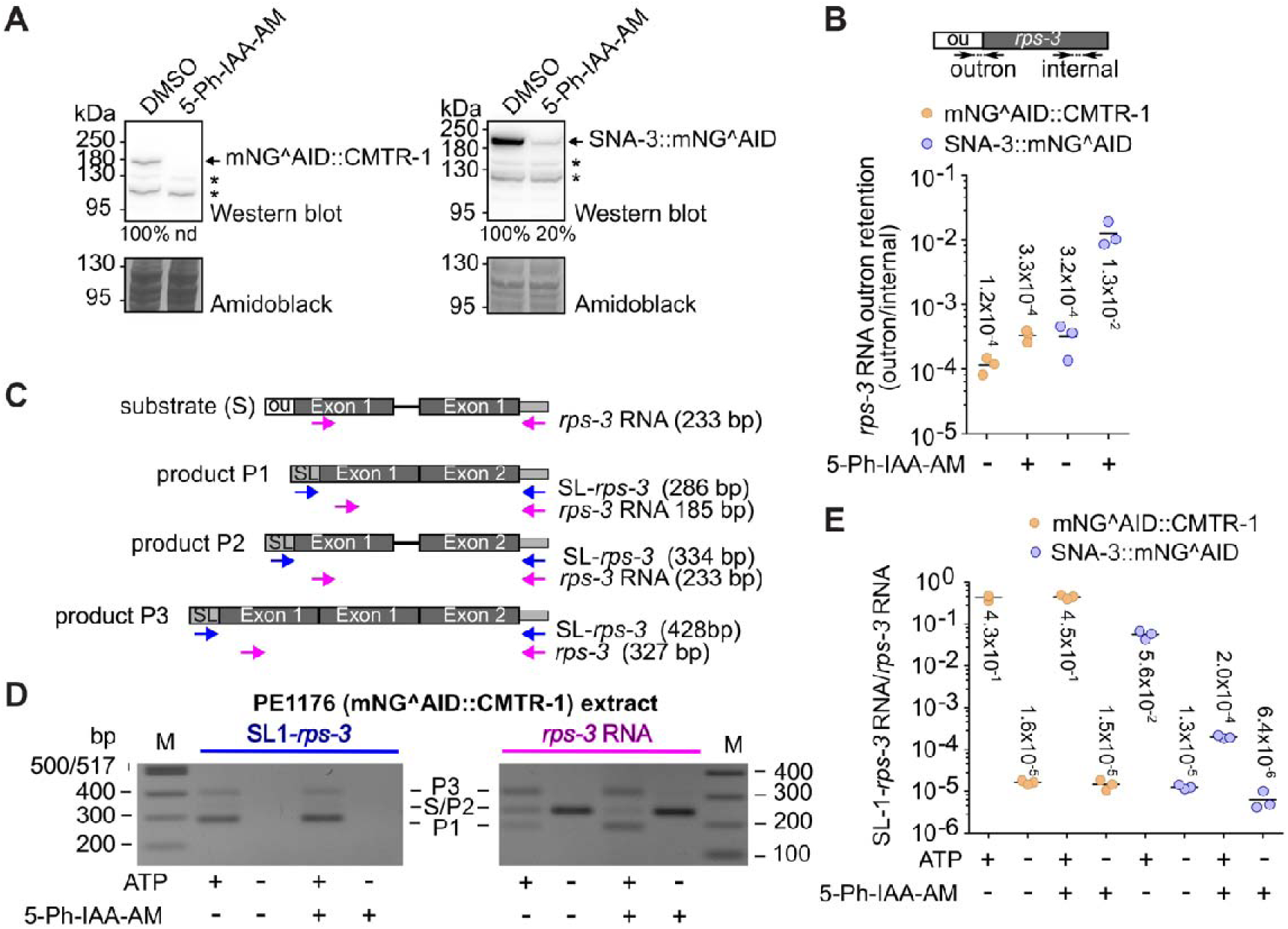
CMTR-1 is not required for spliced leader *trans*-splicing. (A) Western blots showing levels of endogenous mNG^AID::CMTR-1 and SNA-3::mNG^AID proteins in embryonic extracts prepared from PE1176 and PE1220 animals, respectively. Embryos were control-treated with DMSO or with the auxin analogue 5-Ph-IAA-AM for 4h prior to extract preparation. Proteins were analysed by Western blotting and detected by anti- mNeonGreen antibodies. Total protein visualised using amidoblack staining was used to standardise the levels of mNeonGreen-tagged proteins. The level of SNA-3::mNG^AID protein in 5-Ph-IAA-AM treated extract was expressed as percentage of that detected in the DMSO-treated extract. The mNG^AID::CMTR-1 was not detectable in 5-Ph-IAA-AM treated extracts (nd). Non-specific bands are indicated with a star symbol. (B) Quantitative RT-PCR measurement of outron retention of endogenous *rps-3* transcripts. The diagram shows the location of primers that detect outron and internal regions of *rps-3* RNA. RNA was isolated from embryonic extracts, reverse transcribed, and analysed by quantitative PCR as described previously (Philippe et al., 2017). The graph shows the ratio of endogenous *rps-3* outron to internal amplicon levels in PE1176 (gold) and PE1220 (blue) extracts treated with 5-Ph-IAA-AM (+) or control treated with DMSO (-) determined using the ΔC_T_ method (Schmittgen and Livak, 2008). Measurements are technical replicates and their means (horizontal line, mean values are shown). The analysis is representative of 2 independent measurements. (C) Substrate and products of *in vitro* SL *trans*-splicing. Diagrammatic representation of the synthetic *rps-3* transcript (substrate S) with outron (ou) at the 5’ end and the products P1, P2 and P3. The locations of primer pairs SL1-*rps-3* and *rps-3* RNA used for detection by one-step qPCR and expected amplicon size are indicated. (D) Analysis of *in vitro* SL *trans*-splicing by agarose gel electrophoresis. Synthetic *rps-3* RNA was incubated with embryonic extracts from PE1176 embryos treated with 5-Ph-IAA-AM or control-treated with DMSO (+/- 5-Ph-IAA-AM) as described. ATP and ATP regeneration system were included where indicated (+/- ATP). Products were amplified by one-step PCR, analysed by 1.8% agarose gel electrophoresis and visualised by staining with ethidium bromide. (E) Analysis of *in vitro* SL *trans*-splicing by quantitative RT-PCR. Synthetic *rps-3* RNA was incubated with embryonic extracts from PE1176 (gold) or from PE1220 embryos (blue) treated with 5-Ph-IAA-AM or control-treated with DMSO (+/- 5-Ph-IAA-AM) as described. ATP and ATP regeneration system were included where indicated (+/- ATP). Products were quantified by quantitative one-step RT-PCR as described. The graph shows the ratio of *trans*-spliced SL1-*rps-3* products standardised with respect to total synthetic *rps-3* RNA using the ΔC_T_ method (Schmittgen and Livak, 2008). The *in vitro* processing reactions were done in triplicates. Shown are the results for each reaction and the mean (horizontal line, values are indicated).

We analysed the effect of protein depletion on spliced leader *trans*-splicing in embryonic extracts using two assays: 1) the steady state spliced leader *trans*- splicing levels of endogenous *rps-3* transcripts; and 2) the *trans*-splicing by embryonic extracts of an exogenous, synthetic *rps-3* transcript prepared by *in vitro* transcription. In the first assay, depletion of SNA-3 led to a 40.6-fold increase in non- *trans*-spliced, outron-containing endogenous *rps-3* RNAs, while the depletion of CMTR-1 had only a minor effect, leading to a 2.75-fold increase of non-*trans*-spliced *rps-3* RNA (Figure 5B).

SL1 *trans*-splicing of synthetic *rps-3* transcripts was analysed by PCR as previously described (Lasda et al., 2011). The synthetic *rps-3* RNA used as substrate contains plasmid sequence at the 3’-end allowing the use of primers that distinguish the synthetic *rps-3* RNA from endogenous *rps-3* RNA (Figure 5C). Products P1, P2 and P3 (Figure 5C, D) were identified by sequence analysis and are detected by the SL- *rps-3* primers that amplify SL *trans*-spliced synthetic RNA and the *rps-3* RNA control primers that detect all forms of synthetic *rps-3* RNA. Importantly, SL-*rps-3* primer products were only detected in reactions containing ATP necessary for spliced leader *trans*-splicing to occur (Figure 5C). Analysis by qPCR shows that depletion of CMTR- 1 has a minor effect on *in vitro trans*-splicing of the exogenously added synthetic *rps- 3* RNA, with near-identical amounts of product detected in the presence of ATP, independent of whether CMTR-1 was depleted or not (Figure 5D). In contrast, the depletion of SNA-3 led to a 280-fold reduction of detection of *trans*-spliced products (Figure 5E). Thus, both assays failed to detect a clear, pronounced effect of CMTR-1 depletion on spliced leader *trans*-splicing, suggesting that neither cOMe nor CMTR-1 has a significant role in this process.

### Loss of cOMe leads to differential transcript levels for genes involved in the intracellular pathogen response and germline sex determination

To investigate the possible impact of loss of cOMe on gene expression, we compared transcript levels for 5-Ph-IAA treated to control-treated animals. We cultured synchronised populations of mNG^AID::CMTR-1 larvae, and began the 5- Ph-IAA treatment when the animals were at the third larval (L3) stage, corresponding to the developmental stage at which CMTR-1 is critical for the sperm-oocyte switch (Figure 1J). Both control and 5-Ph-IAA treated animals were harvested for RNA preparation at early L4 stage.

We performed differential gene expression analysis, comparing RNA-Seq derived from control versus 5-Ph-IAA treated animals. This revealed 1953 genes whose steady state transcript levels were significantly upregulated by mNG^AID::CMTR-1 depletion (FDR<=0.05, absolute log2 fold change >=1), and 3398 genes for which transcript levels were significantly reduced by the same treatment (FDR<=0.05, absolute log2 fold change >=1) (Supplementary Table 3).

Inspection of the genes that were significantly enriched more than four-fold in mNG^AID::CMTR-1 knock-down worms compared to controls revealed upregulation of around 80 genes, including *fbxa-158*, *pals-14*, and *pals-6*, which belong to a class of genes that constitute the intracellular pathogen response (IPR), a *C. elegans* transcriptional immune response to intracellular pathogens (Lažetić et al., 2023). IPR is induced by viral and microsporidia infections, but also mitochondrial dysfunction(Mao et al., 2022; Sowa et al., 2020).

To systematically investigate the possibility that IPR genes were generally upregulated in response to depletion of CMTR-1, we extracted the fold changes of 272 IPR genes that were upregulated more than four-fold in response to expression of Orsay virus RNA-dependent RNA polymerase (Sowa et al., 2020)(Figure 6A).

**Figure 6.**
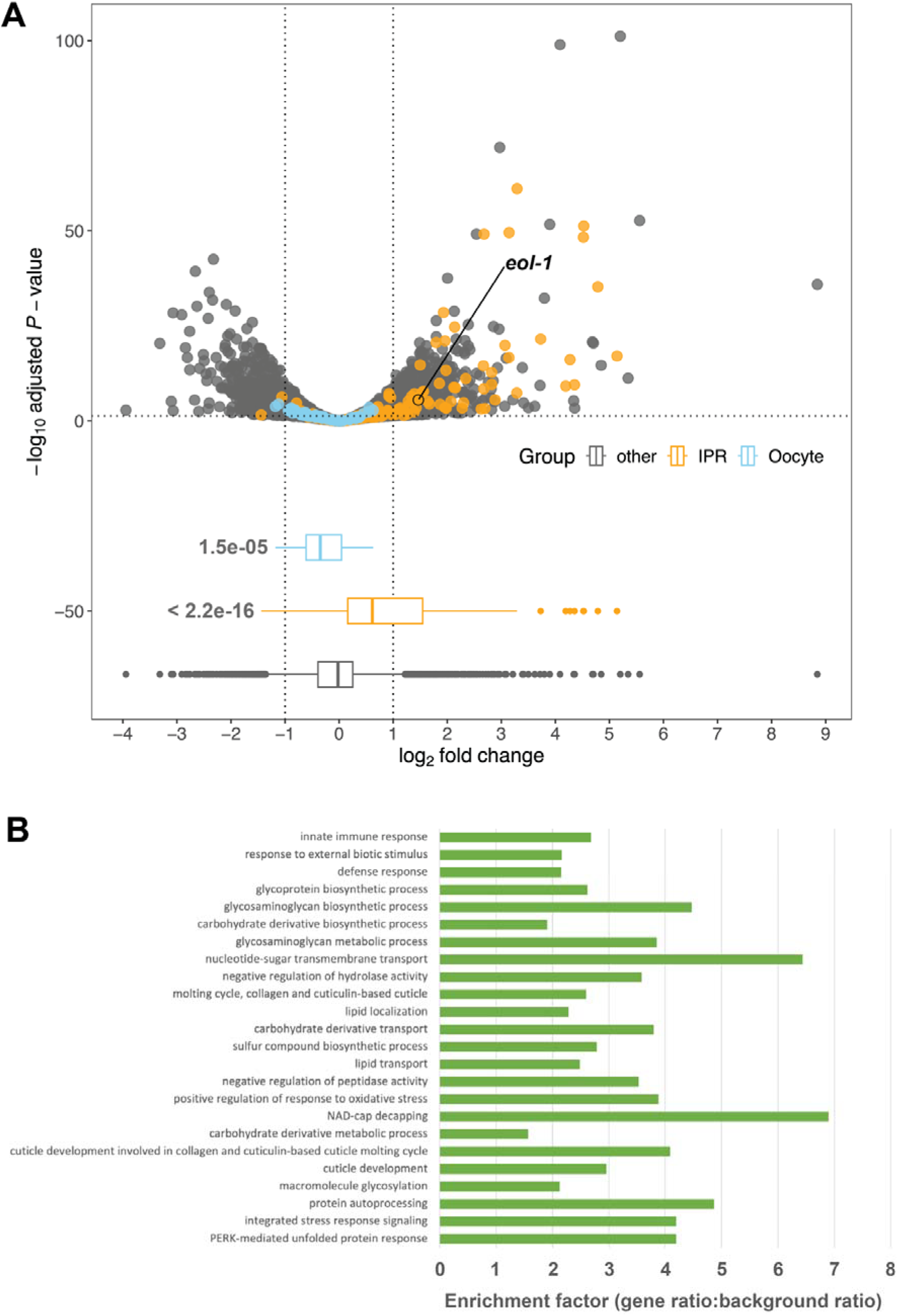
Depletion of CMTR-1 results in upregulation of transcripts associated with the innate immune response and downregulation of transcripts involved in germline sex determination. (A) Upper panel, volcano plot showing differential gene expression in animals subject to CMTR-1 depletion compared to controls (three biological replicates). Intracellular pathogen response genes (IPR) are highlighted in orange. Germline sex determination genes (Oocyte) are highlighted in blue. Lower panel, box plot of the fold change data, showing the p-values (Welch Two Sample t-test) for the differences between the means of the IPR and Oocyte gene sets and the “other” (non-IPR/germline sex determination genes). The gene *eol-1*, which is part of the IPR and was subsequently shown in this study to be epistatic to *cmtr-1* (Figure 7), is also indicated. (B) Gene ontology enrichment analysis showing the significantly enriched gene categories among upregulated genes in animals subject to CMTR-1 depletion compared to controls ranked by q-value.

These IPR genes showed statistically significantly higher transcript levels in response to CMTR-1 depletion compared to non-IPR genes (P<<0.001), though the effect size was modest (mean fold change: 0.697) (Figure 6A). The effect of loss of CMTR-1 on innate immune gene expression was also supported by GeneOntology (GO) term enrichment analysis (Figure 6B): the three most significantly overrepresented GO terms among upregulated genes were “innate immune response” (GO:0045087, qvalue = 2.02x10^-8^), “response to external biotic stimulus” (GO:0043207, qvalue = 3.92x10^-7^), and “defense response” (GO:0006952, qvalue = 4.20x10^-7^). Thus, loss of CMTR-1 and cOMe leads to transcriptional upregulation of many intracellular pathogen response genes.

GO term enrichment analysis among downregulated genes in response to loss of CMTR-1 highlighted a broad range of metabolic terms but, intriguingly, also terms related to gametogenesis, including “gamete generation” (GO:0007276, qvalue = 3.59 x10^-13^) and “germ cell development” (GO:0007281, qvalue = 1.96x10^-9^) (Supplementary Table 3).

To investigate whether transcripts from genes associated with germline sex determination were downregulated in response to depletion of CMTR-1, we extracted the fold changes of a set of 64 genes involved in germline sex determination (Figure 6A). We found that this set of genes showed statistically significantly (P<<0.001) lower fold changes than all other genes, though as for the IPR gene response, the effect size was modest (mean fold change: -0.287).

Nonetheless, since germline development is known to be highly sensitive to shifts in gene expression (Ellis, 2022), this overall trend of downregulation may explain the germline defects that we observed (Figure 1).

We were unable to establish a clear correlation between transcript level changes and the growth defects that we observe in animals depleted of CMTR-1. It is likely that changes in the expression of a broad range of genes are responsible for this phenotype.

### Loss of the RNA decapping exonuclease EOL-1 bypasses the need for CMTR-1

To better understand the molecular and cellular significance of the global loss of cOMe, we performed an unbiased screen for suppressors of the *cmtr-1* loss-of-function phenotype, reasoning that such mutations might allow us to identify components involved in the cellular response to cOMe modified RNAs.

We mutagenised PE1176 worms with ethyl methanesulfonate (EMS) and grew the F2 generation on NGM plates containing 1 µM 5-Ph-IAA. We then screened for the presence of healthy gravid worms, which would indicate suppression of the *cmtr-1* loss-of-function phenotype. Suppression was confirmed by showing that the worms could be maintained over multiple generations on 5-Ph-IAA containing media. Only one strain was retained per F1 culture, to ensure independence of each suppressor mutation. From combined screens of approximately 20,000 haploid genomes, we recovered 27 independent mutants that were fertile when grown in the presence of 5-Ph-IAA.

We designed the screen with the expectation that we would recover loss of function mutations in the *TIR1(F79G)* transgene, which restore fertility simply by preventing 5-Ph-IAA-depdenent degradation of mNG^AID ::CMTR-1. Such mutations, which constitute a type of informational suppression (Mount and Anderson, 2000), could be distinguished from *bona fide cmtr-1* loss-of-function suppressors by the fact that they express mNG::AID^CMTR-1 in the presence of 5-Ph-IAA (Supplementary Figure 2). Using this secondary screen, we found that of the 27 mutants, 21 showed nuclear mNeonGreen fluorescence despite being grown on 5-Ph-IAA. We sequenced the TIR1 coding region from six of the putative TIR1 loss-of-function mutants and confirmed that all were missense mutations (Supplementary Figure 2).

The six mutants that lacked mNeonGreen fluorescence but were nonetheless fertile were thus putative suppressors that bypass the requirement for CMTR-1. We confirmed that they also suppressed the *cmtr-1(syb3613)* null allele: all six mutations suppressed the sterile phenotype and growth defects conferred by the *syb3613* allele confirming that they are epistatic to *cmtr-1* loss-of-function (Figure 7A).

**Figure 7.**
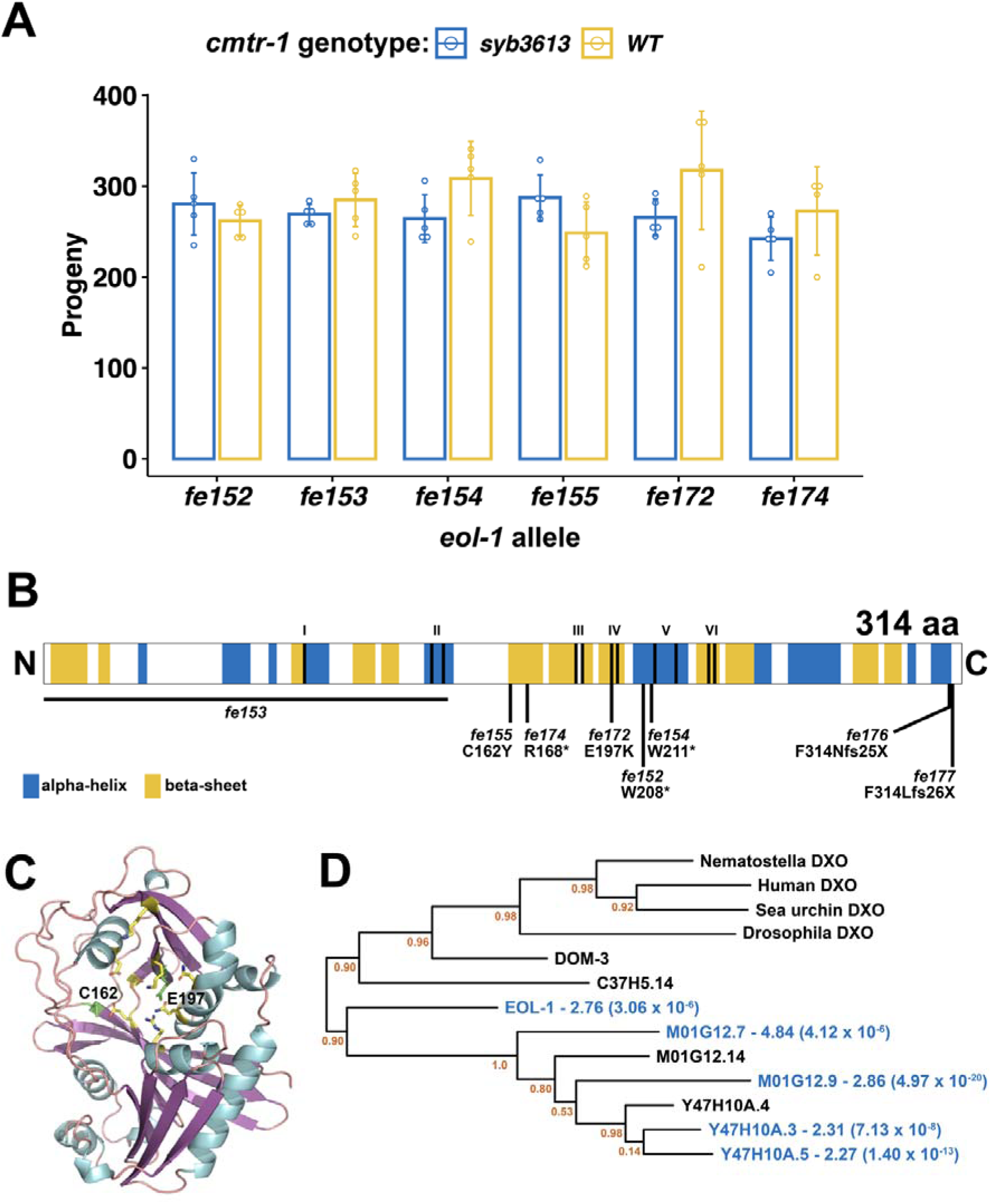
Loss of function mutations in *eol-1* suppress the loss of *cmtr-1* mutant phenotype. (A) Brood counts for the six *eol-1* alleles identified as suppressors of *cmtr-1* loss-of-function in *cmtr-1(syb3613)* and *cmtr-1(+)* backgrounds. Histograms show means and error bars show standard deviations for counts derived from five adults for each genotype. (B) Schematic of EOL-1 showing the location of the suppressor mutations and the two mutations generated by Cas9/CRISPR (*fe176* and *fe177*). Shading indicates secondary structure assignments based on the AlphaFold prediction (A5JYX9). Roman numerals indicate highly conserved motifs necessary for decapping and exonuclease activity (Wang et al., 2015). (C) AlphaFold prediction for EOL-1, showing location of residues affected by missense suppressor mutations (green). Active site residues other than E197 are in yellow. (D) Phylogram showing the relationship between EOL-1 and other DXO family members. Blue text indicates proteins with transcripts that are significantly enriched in mNG^AID::CMTR- 1 depleted animals. Fold enrichment is given, along with adjusted *p* values in brackets. Node statistical supports are given as aLRT (SH-like) values.

To identify the gene(s) affected by these six mutations, we used a whole-genome sequencing (WGS) mapping strategy based on the sibling-subtraction approach described previously (Joseph et al., 2017). We mapped two of the suppressor mutants, *fe152* and *fe154*, using this approach (Supplementary Figure 3). The sibling-subtraction WGS mapping approach relies on the identification of variants that are present in more than 90% of reads derived from the pooled broods of F2 suppressor animals, but absent, or at a low frequency in the reads derived from their non-suppressor sibling broods. In practice, we found that focussing only on variants found in 100% of suppressor pool reads was sufficient to narrow down the location of the suppressor mutations to a region of chromosome V (Supplementary Table 4).

SnpEff and manual curation of these variants revealed predicted damaging variants in a single gene, *eol-1*, allowing us to assign *fe152* and *fe154* as premature stop codon mutations, W208Opal and W211Amber, respectively (Figure 7B).

Sanger sequencing of the *eol-1* locus in the other four suppressor mutants revealed that each was a predicted damaging allele of *eol-1* (Figure 7B). A third nonsense mutation is created by *fe174*, R168Opal, and two mutations, *fe155* and *fe172*, are missense mutations, C162Y and E297K, respectively. The *fe153* allele was found to be a deletion of 1456 bp that removes 940 bp upstream of the start codon as well as half of the coding region. The deletion is accompanied by insertion of 349 bp of sequence, part of which appears to be repetitive DNA found on multiple *C. elegans* chromosomes.

Having identified the genetic location of the suppressor mutations, we crossed them into a *cmtr-1* wild-type background, which revealed that they had wild-type brood sizes (Figure 7A). Thus, loss of *eol-1* function is compatible with wild-type fertility. We also did not detect any obvious phenotypes that would indicate defects in embryonic or postembryonic development.

Although the recovery of six independent function altering mutations in the same gene is strong evidence that loss of *eol-1* function suppresses the requirement for CMTR-1 and thus cOMe, we wanted to formally confirm this. We aimed to delete the *eol-1* gene, using Cas9/CRISPR to generate double-strand breaks at either end of the gene. However, we found that the guide RNA targeted to the 5’-end of the coding region was ineffective, so we were not able to identify gene deletions. Nonetheless, we identified two alleles, *fe176* and *fe177*, that created frameshifts just upstream of the stop codon, both of which would be predicted to result in peptide extensions that derive from translation into the 3’-UTR (Figure 7B). Proteins with such C-terminal extensions are subject to “readthrough mitigation” mechanisms that lead to downregulation of the affected protein (Arribere et al., 2016; Kesner et al., 2023; Müller et al., 2023). Consistent with this, both alleles suppress the sterile phenotype caused by depletion of CMTR-1, confirming that loss of *eol-1* function bypasses the requirement for cOMe.

EOL-1 is a member of the DXO/Rai1 family of decapping enzymes (Chang et al., 2012; Jiao et al., 2013; Mao et al., 2020; Shen et al., 2014), and some members are also distributive 5’ to 3’ exonucleases, so that they can both decap and degrade

their target RNAs. The missense mutations, C162Y and E297K, alter residues that are part of the predicted active site of EOL-1 (Figure 7C). Indeed, E297 is the homologue of mammalian DXO E253, a critical catalytic residue, part of the so-called Motif IV (Chang et al., 2012; Jiao et al., 2013). Thus EOL-1 loss of function mutations which specifically abolish its decapping/exonuclease activity, completely suppress the sterility caused by loss of CMTR-1, indicating that removing this activity is likely the mechanism which mediates the suppression.

Previous studies have shown that EOL-1 is a prominent part of the intracellular pathogen response. Given that loss of CMTR-1 led to enrichment of transcripts associated with this response, we looked for evidence that *eol-1* transcripts are enriched in CMTR-1 depleted animals compared to controls (Figure 6A) and found that they are enriched 2.76-fold. EOL-1 is one of nine *C. elegans* DXO homologues (Mao et al., 2020), and is part of a clade of paralogues (Figure 7D). Transcripts from four of these are also upregulated by depletion of CMTR-1, though only two of them are broadly upregulated as part of the intracellular pathogen response (Supplementary Table 3).

## DISCUSSION

Insights into the role of cOMe in animal development requires that one study the effects of loss of the CMTR proteins in the context of whole organism biology. While work in mice and flies have advanced our understanding of cOMe biology, the fact that these RNA modifications are essential for mammalian embryogenesis, but dispensable in flies and some cultured mammalian cells is puzzling. By investigating cOMe function in *C. elegans*, we have obtained a broader evolutionary perspective of the role of these RNA modifications.

Unlike the situation in mammals (Furuichi et al., 1975; Werner et al., 2011), we find that CMTR-1 is the principal enzyme involved in adding cOMe to *C. elegans* RNAs, and consistent with this, has higher steady state nuclear levels compared to CMTR-2. Similarly consistent with these observations, CMTR-1 is essential while CMTR-2 is not. We did not find a function for CMTR-2 beyond augmenting that of CMTR-1, which suggests that CMTR-2 is also able to modify the first nucleotide, a substrate specificity also found for its *Drosophila* and human homologues in modifying canonical AGU starting transcripts (Anreiter et al., 2020; Dix et al., 2022; Haussmann et al., 2022).

The two nematode CMTR proteins have distinct sub-cellular localisations and interaction partners, suggesting that they are functionally differentiated. Much of this may be due to their differing subcellular localisations. As in mammals and *Drosophila* (Haussmann et al., 2022; Werner et al., 2011), we see stronger associations of CMTR-2 with the cytoplasm. This is most pronounced during embryogenesis where CMTR-2 is apparently excluded from the nucleus, something not reported in other animal cells. This observation warrants further investigation.

The protein interaction profile of *C. elegans* CMTR-1 is highly similar to its mammalian homologue (Simabuco et al., 2019), with both proteins showing prominent interactions with the spliceosome. The functional significance of these observations is unclear but may simply reflect the fact that these proteins assemble on nascent transcripts, though we cannot rule out a functional role for CMTR-1 in *cis*- splicing of certain transcripts. However, our experiments do indicate that CMTR-1 is not an essential component of nematode spliced leader *trans*-splicing. Despite the interaction between CMTR-1 and SNA-3, neither of our functional assays indicates that CMTR-1 is required for the activity of the *trans*-spliceosome.

### A role for EOL-1 in degrading RNAs lacking cap-adjacent 2’-O-ribose methylation

Since cOMe plays an essential physiological role in *C. elegans*, this has allowed us to fully exploit the advantages of this genetically tractable model system to understand the mechanistic basis of the cellular response to cOMe. Importantly, we have identified EOL-1 as a key effector that recognises transcripts lacking cOMe. A previous study showed that cOMe blocks the activity of mammalian DXO *in vitro* (Picard-Jean et al., 2018). The genetic interaction between *cmtr-1* and *eol-1* demonstrates that this interaction is physiologically significant. Since we find that loss of EOL-1 effectively bypasses the requirement for CMTR-1, and hence cOMe, our work provides clear support for the hypothesis that DXO proteins recognise and destroy RNAs that lack cOMe (Figure 8). Thus, in animals lacking CMTR-1, all cellular transcripts become potential targets, leading to changes in steady-state transcript levels that we see correlated with defects in germline development.

**Figure 8.**
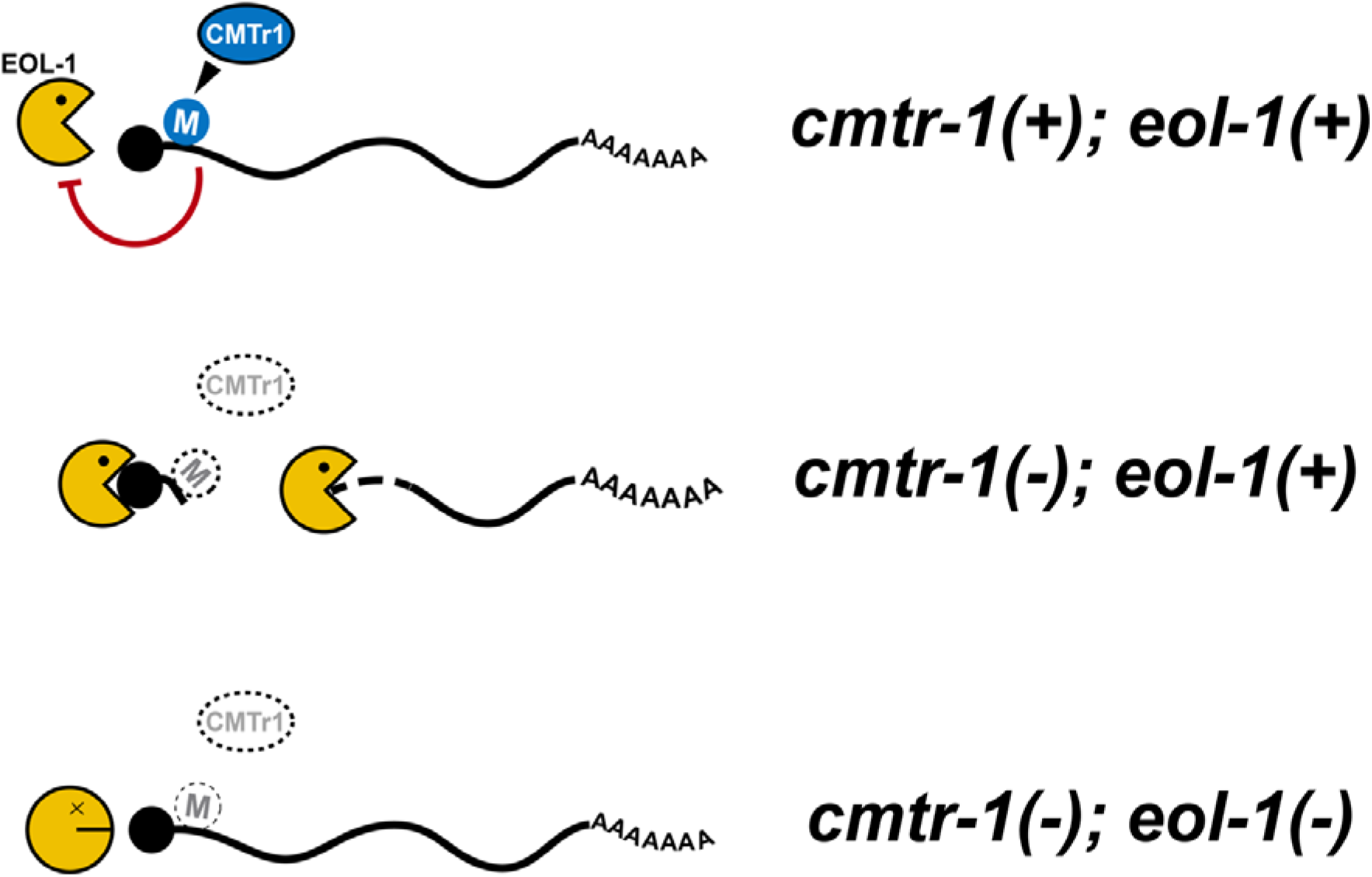
Cap-adjacent 2’-O-ribose methylation protects cellular transcripts from EOL-1-mediated degradation. In this proposed model, EOL-1 is prevented from decapping cOMe modified transcripts (M). In the absence of CMTR-1, transcripts lack cOMe and thus become substrates for EOL-1. Transcripts for genes that are sensitive to copy number changes, such as those involved in germline cell fate, would be most susceptible to loss of cOMe. Loss of EOL-1 prevents the degradation of these transcripts. Black shaded circle indicates the monomethyl guanosine cap.

What is the physiological significance of EOL-1 activity? In wild-type animals, most, if not all, RNA polymerase II transcripts will possess this modification and thus would be protected from EOL-1 degradation. Previous studies have proposed that DXO proteins maintain RNA quality control, removing partially capped RNAs that lack cOMe before they enter the cytoplasm (Jiao et al., 2013). They have also been shown to have a high affinity for NAD-capped RNAs, which are targeted by DXO for rapid degradation (Kiledjian, 2018). Such roles would be consistent with our data.

However, since EOL-1 is a prominent component of the intracellular pathogen response, a role in innate immunity is also plausible; indeed, previous studies of this protein show that it is intimately associated with responses to pathogens and cellular dysfunction.

EOL-1 was first identified in screens for mutants with enhanced aversive behavioural responses to pathogenic bacteria (Shen et al., 2014), and it is a notable member of the intracellular pathogen response, induced by Orsay virus, a positive sense, single-stranded RNA virus, and the microsporidian *Nematocida parisii* (Chen et al., 2017; Tecle et al., 2021). Mutations associated with mitochondrial dysfunction also result in elevated *eol-1* expression levels (Mao et al., 2020; Senchuk et al., 2018), and EOL-1 protein undergoes changes in its intracellular localisation, forming cytoplasmic foci in response to mitochondrial defects (Mao et al., 2020). However, a mechanistic role for EOL-1 in innate immunity has yet to be defined. Our work connecting EOL-1 to cOMe suggests a possible function for this protein in the surveillance for RNAs lacking cap1 structures. Mitochondrial dysfunction leads to the presence of RNAs in the cytoplasm that lack cOMe (Mao et al., 2020), which would presumably be substrates for EOL-1 degradation. Moreover, mitochondrial RNAs also possess high levels of NAD-caps, making them preferential DXO family substrates (Bird et al., 2018; Jiao et al., 2017). Likewise, Orsay virus RNAs, for instance, lack both a 5’ monomethyl guanosine cap and cOMe modifications (Ashe et al., 2013). Thus, the identification and removal of pathogenic RNAs on the basis of cOMe absence is a plausible function of this enzyme in *C. elegans* that warrants further exploration.

The relationship we have discovered between cOMe and EOL-1 may have broader phylogenetic significance, with DXO proteins having a conserved role in RNA surveillance based on cOMe status. In particular, this may explain the molecular basis of the lethality observed in mammalian embryos lacking either of the two CMTRs (Dohnalkova et al., 2023); in the light of our work, perhaps loss of cOMe in mouse embryos renders many cellular transcripts targets of mammalian DXO. While *DXO* transcripts are not upregulated by the loss of cOMe (Dohnalkova et al., 2023), unlike those of *eol-1* in *C. elegans*, cellular levels of DXO may still be sufficient to have an impact across the transcriptome. Thus, the embryonic lethality observed in embryos lacking cOMe might be explained by the cumulative downregulation of multiple transcripts. It will be important to determine whether loss of DXO can suppress the embryonic lethality of loss of *Cmtr1* in mice.

### Cap-adjacent 2’-O-ribose methylation and the innate immune response

Our findings highlight similarities between the functions of cOMe in nematodes and mammals: as in mammals, loss of cOMe leads to the upregulation of genes associated with an innate transcriptional immune response. This raises the possibility that cOMe is a determinant that regulates an RNA-mediated immune response in *C. elegans*. The upregulation of IPR genes that we observed in animals depleted of CMTR-1 is intriguing given that loss of CMTR1 in mammals also results in induction of the innate immune response (Daffis et al., 2010; Dohnalkova et al., 2023; Schuberth-Wagner et al., 2015). In mammals, RNAs lacking cOMe are directly sensed by RIG-I, which binds to double-stranded RNA substrates that lack cOMe and leads to the activation of the type I interferon response (Devarkar et al., 2016; Schuberth-Wagner et al., 2015); although the mechanism by which cellular RNAs are converted into double-stranded RNAs remains unclear.

In *C. elegans*, there are three RIG-I homologues, DRH-1, DRH-2 and DRH-3 (Guo et al., 2013), but DRH-1 appears to be functionally most similar to RIG-I in that it recognises viral RNAs and is essential for the intracellular pathogen response to viral infection (Sowa et al., 2020). Indeed, the double-stranded RNA sensing module (the helicase and C-terminal domains) of RIG-I can functionally substitute for the homologous region of DRH-1 in assays for viral load (Guo et al., 2013); and native DRH-1 binds to double-stranded RNAs in a manner similar to RIG-I (Consalvo et al., 2024). Thus, if the upregulation of the intracellular pathogen response is a direct effect of appearance of RNAs lacking cOMe, then DRH-1is a good candidate involved in their detection; it will be important to test whether loss of *drh-1* results in abrogation of the intracellular pathogen response in animals lacking cOMe.

An alternative possibility is that the induction of the intracellular pathogen response by loss of cOMe is indirect, caused by the overall impact that this loss has on global gene expression patterns. The intracellular pathogen response is also induced by proteotoxic stress (Bakowski et al., 2014; Panek et al., 2020; Reddy et al., 2017), so the global, cumulative effects on gene expression caused by loss of cOMe, might lead to an intracellular pathogen response due to disruption of proteostasis.

Distinguishing between direct and indirect effects of cOMe on the transcriptional immune response in *C. elegans* will be important in determining the extent of functional conservation of cOMe function between nematodes and mammals.

Multiple studies in diverse experimental systems have described roles for cOMe at multiple steps in the regulation of gene expression, including transcription, splicing and translation (Dönmez et al., 2004; Drazkowska et al., 2022; Haussmann et al., 2022; Kuge et al., 1998; Liang et al., 2022; Yu et al., 1998). It is thus surprising that in *C. elegans* that lack EOL-1, cOMe is largely dispensable, at least in laboratory culture. However, our results are consistent with work showing that in certain mammalian cell types, the impact of loss of cOMe is relatively minor (Dohnalkova et al., 2023). Similarly, loss of cOMe in *Drosophila* does not result in profound cellular impacts, with such animals being viable and fertile (Haussmann et al., 2022). The ubiquity of cOMe and the conservation of the two CMTR proteins that catalyse it shows that it must have an essential role across animal phylogeny. Perhaps its original, ancestral role was in RNA-based immune surveillance, which has been preserved in *C. elegans* and mammals (but apparently lost in *Drosophila*), with its functions in gene expression having arisen due cellular adaptation to the presence of cOMe on cellular RNAs. Alternatively, it may have been co-opted into roles in innate immunity through convergent evolution in mammals and nematodes from an ancestral function(s) in the regulation of gene expression. Better understanding of the molecular impacts of cOMe loss in diverse physiologically relevant experimental systems will be needed to distinguish these hypotheses.

## DATA AVAILABILITY

The mass spectrometry proteomics data have been deposited to the ProteomeXchange Consortium via the PRIDE (Perez-Riverol et al., 2022) partner repository with the dataset identifier PXD058991 (See also Supplementary Table 2). The RNA-Seq data used for the CMTR-1 depletion differential gene expression analysis have been deposited in ArrayExpress under accession number E-MTAB- 14816. Sequence reads that were used in the sibling selection whole genome sequence mapping of *eol-1(fe152)* and *eol-1(fe154)* are available in the NCBI Sequence Read Archive under BioProject ID PRJNA1216994.

## SUPPLEMENTARY DATA STATEMENT

Supplementary Data are available at NAR Online.

## ACKNOWLEDGEMENTS

We thank Omar Alotaibi, Holly Armstrong, Sandeep Goli, Phurichaya Khunthong, and Ursula Terech Rial for their assistance with the *cmtr-1* suppressor screen. We also thank Max Bracken-Zmuda for help making the *cmtr-1(feK244A)* strain. Some strains were provided by the Caenorhabditis Genetics Center, which is funded by NIH Office of Research Infrastructure Programs (P40 OD010440). Proteomic analysis was performed by Aberdeen Proteomics, and we thank Kate Burgoyne and Craig Pattinson for technical support. We thank WormBase for providing the community resource that facilitated the interrogation of *C. elegans* molecular genetics used in this work. We acknowledge the support of the Maxwell HPC computer cluster funded by the University of Aberdeen.

## AUTHOR CONTRIBUTIONS

J.P., B.M. and M.S. conceived the research and provided oversight and leadership responsibility for the research activity planning and execution. All authors were involved in performing experiments and data collection. M.W., B.M. and J.P. designed and implemented the computational analyses. E.C., P.E., M.K., M.W., I.H., M.S., B.M. and J.P. prepared figures and tables. E.C., P.E., B.M. and J.P. wrote the manuscript, and all authors reviewed and edited the final manuscript.

## FUNDING

This work was supported by the UKRI Biotechnology and Biological Sciences Research Council (BBSRC) through the EastBio Doctoral Training Partnership (BB/T00875X/1) and a project grant awarded to M.S. (BB/X008193/1).

## SUPPLEMENTARY FIGURES

**Supplementary Figure 1.**
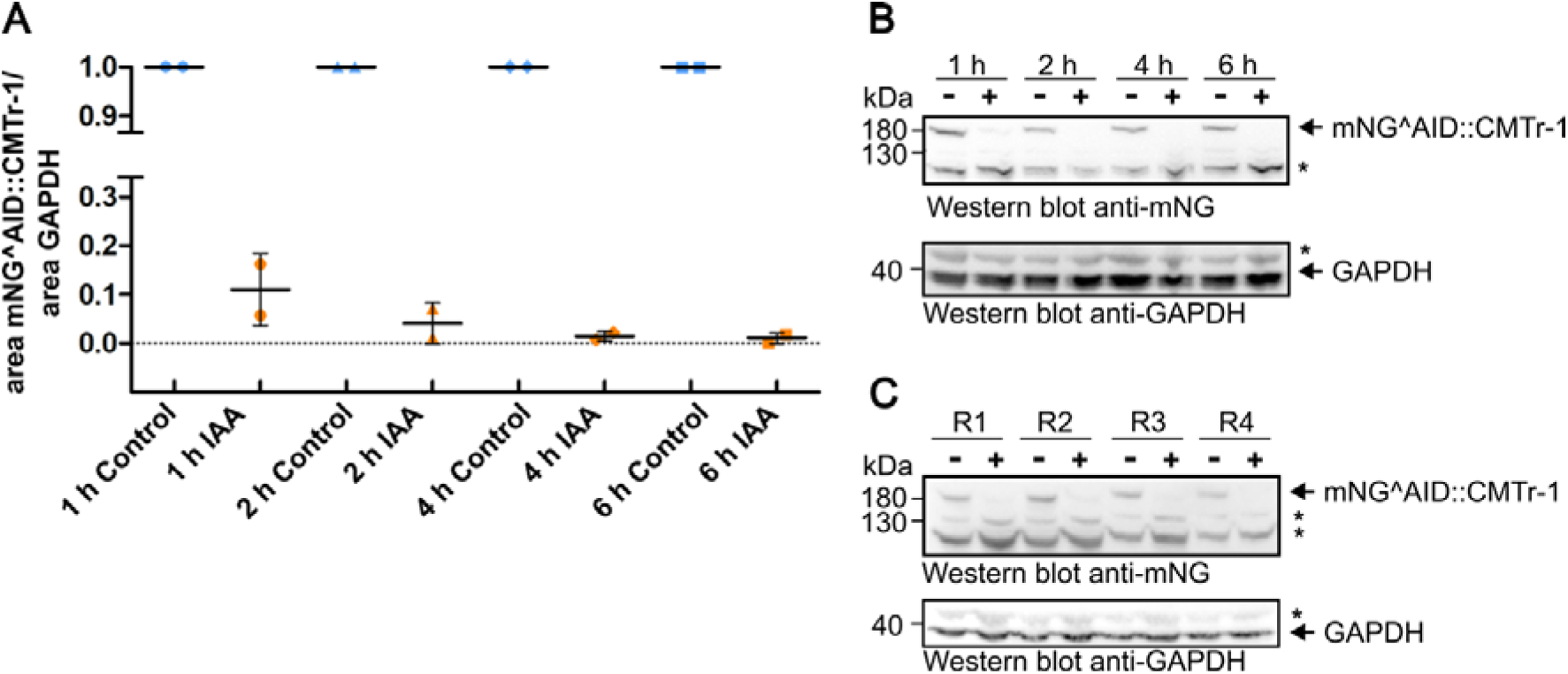
5-Ph-IAA depletion of mNG^AID::CMTR-1. (A) PE1176 animals were treated with 5-Ph-IAA or control-treated and subjected to Western blotting at the indicated time points. Proteins were detected using anti-NeonGreen and anti-GAPDH antibodies. Protein levels of mNG^AID::CMTR-1 were standardised relative to GADPH, with the controls set to 1. Data represents 2 biological replicates. (B) Representative Western blot for the data shown in A. (C) Western blot of mNG^AID::CMTR-1 depletion of replicates (R1- 4) used to generate RNA-Seq data (see Figure 6). In B and C, “+” indicates 5-Ph-IAA treated samples and “-“ control samples, respectively.

**Supplementary Figure 2.**
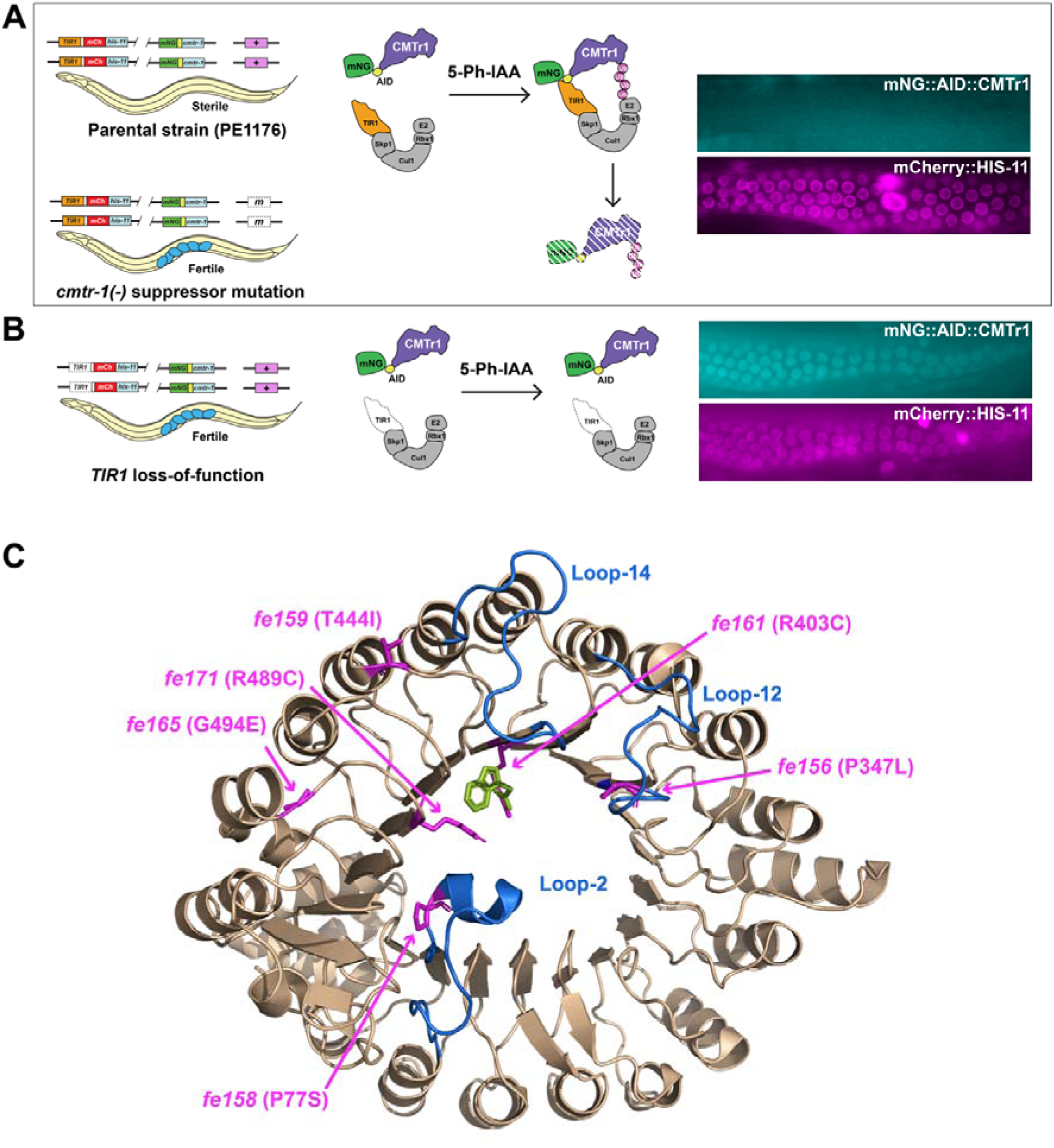
Secondary screen to distinguish suppressors epistatic to *cmtr-1(-)* from *TIR1* loss-of-function mutants. (A) The starting strain for the suppressor mutagenesis screen (PE1176) is sterile, while strains carrying suppressor mutations (m, unshaded, dashed box) that are epistatic to *cmtr-1(-)* restore fertility (blue ovals indicate eggs inside gravid hermaphrodites). (B) We predicted that TIR1 loss-of-function mutants (unshaded, dashed box) would also restore fertility to *cmtr-1(-)* animals, since these would prevent depletion of mNG^AID::CMTR-1. To distinguish between these two suppressor classes, we conducted a secondary screen based on mNeonGreen fluorescence. This was absent in PE1176 and epistatic suppressor strains (panels show distal gonad arms of adult hermaphrodites; nuclear mCherry::HIS-11 fluorescence was unaffected and serves to locate the germline nuclei), but was present in *TIR1* loss-of-function mutants. (C) Location of selected *TIR1* loss-of-function mutations identified from secondary screening (magenta). View of the TIR1-LRR domain showing the auxin and substrate-binding pocket (https://doi.org/10.2210/pdb2P1Q/pdb; (Tan et al., 2007)). Auxin is shown in green, the three extended loops key for the formation of the pocket are shown in blue.

**Supplementary Figure 3.**
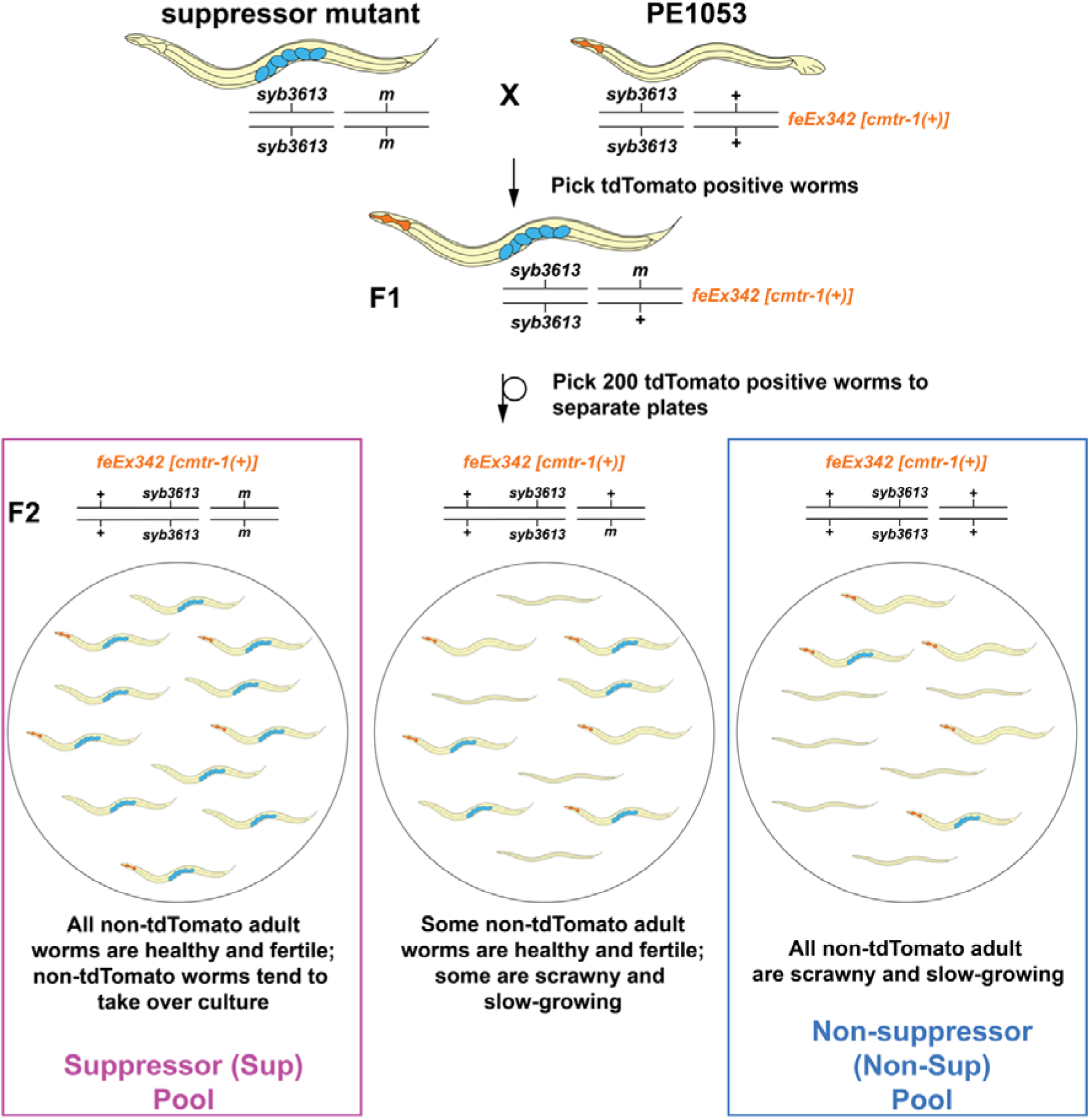
Strategy used to isolate strains for sibling-subtraction/whole genome sequencing mapping. Suppressor strains were crossed to the transgenic rescue line, PE1053, which is homozygous for *cmtr-1(syb3613)* but rescued by the *feEx342 cmtr-1(+)* transgene. The Sup and Non-Sup lines were established from single F2 animals on the basis that F2 suppressor homozygotes were not dependent on the *feEx342 cmtr-1(+)* transgene, while the broods of non-suppressor homozygotes resembled those of the PE1053 grandparent strain. Note, *feEx342* shows relatively poor rescue of the *cmtr-1* loss-of-function phenotype, so some transgenic animals are sterile. Transgenic animals were recognised on the basis of the *myo- 2p::tdTomato* expression in the pharynx, which is also carried on the *feEx342* extrachromosomal transgenic array. Blue ovals indicate eggs inside gravid hermaphrodites.

## SUPPLEMENTARY TABLES

**Supplementary Table 1.**

Strains and oligonucleotides.

**Supplementary Table 2.**

GFP::CMTR-1 and GFP::CMTR-2 immunoprecipitation data.

**Supplementary Table 3.**

PE1176 5-Ph-IAA treated versus control differential gene expression and GO enrichment analysis.

**Supplementary Table 4.**

Variant mapping for *fe152* and *fe154*. Only variants found in 100% of Sup reads, and absent in 100% of Non-Sup reads are shown. Highlights indicate variants found in both *fe152* and *fe154* Sup pools.

## Notes

### Competing Interest Statement

The authors have declared no competing interest.

### Summary of Updates

Abstract updated to correct typographical error.

## REFERENCES

Adams JM, Cory S. 1975. Modified nucleosides and bizarre 5’-termini in mouse myeloma mRNA. Nature 255:28–33.

Andrews S. 2010. FastQC: a quality control tool for high throughput sequence data. https://www.bioinformatics.babraham.ac.uk/projects/fastqc/

Anreiter I, Mir Q, Simpson JT, Janga SC, Soller M. 2020. New Twists in Detecting mRNA Modification Dynamics. Trends Biotechnol. doi:10.1016/j.tibtech.2020.06.002

Anreiter I, Tian YW, Soller M. 2023. The cap epitranscriptome: Early directions to a complex life as mRNA. Bioessays 45:e2200198.

Arribere JA, Bell RT, Fu BXH, Artiles KL, Hartman PS, Fire AZ. 2014. Efficient marker-free recovery of custom genetic modifications with CRISPR/Cas9 in *Caenorhabditis elegans*. Genetics 198:837–846.

Arribere JA, Cenik ES, Jain N, Hess GT, Lee CH, Bassik MC, Fire AZ. 2016. Translation readthrough mitigation. Nature 534:719–723.

Ashburner M, Ball CA, Blake JA, Botstein D, Butler H, Cherry JM, Davis AP, Dolinski K, Dwight SS, Eppig JT, Harris MA, Hill DP, Issel-Tarver L, Kasarskis A, Lewis S, Matese JC, Richardson JE, Ringwald M, Rubin GM, Sherlock G. 2000. Gene ontology: tool for the unification of biology. The Gene Ontology Consortium. Nat Genet 25:25–29.

Ashe A, Bélicard T, Le Pen J, Sarkies P, Frézal L, Lehrbach N, Félix M, Miska E. 2013. A deletion polymorphism in the *Caenorhabditis elegans* RIG-I homolog disables viral RNA dicing and antiviral immunity. Elife 2. doi:10.7554/eLife.00994

Bakowski MA, Desjardins CA, Smelkinson MG, Dunbar TL, Lopez-Moyado IF, Rifkin SA, Cuomo CA, Troemel ER. 2014. Ubiquitin-mediated response to microsporidia and virus infection in *C. elegans*. PLoS Pathog 10:e1004200.

Bélanger F, Stepinski J, Darzynkiewicz E, Pelletier J. 2010. Characterization of hMTr1, a human Cap1 2’-O-ribose methyltransferase. J Biol Chem 285:33037–33044.

Benjamini Y, Hochberg Y. 1995. Controlling the false discovery rate: A practical and powerful approach to multiple testing. J R Stat Soc Series B Stat Methodol 57:289–300.

Bird JG, Basu U, Kuster D, Ramachandran A, Grudzien-Nogalska E, Towheed A, Wallace DC, Kiledjian M, Temiakov D, Patel SS, Ebright RH, Nickels BE. 2018. Highly efficient 5’ capping of mitochondrial RNA with NAD+ and NADH by yeast and human mitochondrial RNA polymerase. Elife 7. doi:10.7554/eLife.42179

Chang JH, Jiao X, Chiba K, Oh C, Martin CE, Kiledjian M, Tong L. 2012. Dxo1 is a new type of eukaryotic enzyme with both decapping and 5’-3’ exoribonuclease activity. Nat Struct Mol Biol 19:1011–1017.

Chen K, Franz CJ, Jiang H, Jiang Y, Wang D. 2017. An evolutionarily conserved transcriptional response to viral infection in Caenorhabditis nematodes. BMC Genomics 18:303.

Cingolani P, Platts A, Wang LL, Coon M, Nguyen T, Wang L, Land SJ, Lu X, Ruden DM. 2012. A program for annotating and predicting the effects of single nucleotide polymorphisms, SnpEff. Fly (Austin) 6:80–92.

Consalvo CD, Aderounmu AM, Donelick HM, Aruscavage PJ, Eckert DM, Shen PS, Bass BL. 2024. *Caenorhabditis elegans* Dicer acts with the RIG-I-like helicase DRH-1 and RDE-4 to cleave dsRNA. Elife 13. doi:10.7554/eLife.93979

Daffis S, Szretter KJ, Schriewer J, Li J, Youn S, Errett J, Lin T-Y, Schneller S, Zust R, Dong H, Thiel V, Sen GC, Fensterl V, Klimstra WB, Pierson TC, Buller RM, Gale M Jr, Shi P-Y, Diamond MS. 2010. 2’-O methylation of the viral mRNA cap evades host restriction by IFIT family members. Nature 468:452–456.

Danecek P, Bonfield JK, Liddle J, Marshall J, Ohan V, Pollard MO, Whitwham A, Keane T, McCarthy SA, Davies RM, Li H. 2021. Twelve years of SAMtools and BCFtools. Gigascience 10:giab008.

DeMott E, Dickinson D, Doonan R. 2021. Highly improved cloning efficiency for plasmid-based CRISPR knock-in in *C. elegans*. MicroPubl Biol 2021. doi:10.17912/micropub.biology.000499

Despic V, Jaffrey SR. 2023. mRNA ageing shapes the Cap2 methylome in mammalian mRNA. Nature 614:358–366.

Devarkar SC, Wang C, Miller MT, Ramanathan A, Jiang F, Khan AG, Patel SS, Marcotrigiano J. 2016. Structural basis for m7G recognition and 2’-O-methyl discrimination in capped RNAs by the innate immune receptor RIG-I. Proc Natl Acad Sci U S A 113:596–601.

Dickinson DJ, Pani AM, Heppert JK, Higgins CD, Goldstein B. 2015. Streamlined Genome Engineering with a Self-Excising Drug Selection Cassette. Genetics 200:1035–1049.

Dix TC, Haussmann IU, Brivio S, Nallasivan MP, HadzHiev Y, Müller F, Müller B, Pettitt J, Soller M. 2022. CMTr mediated 2’-O-ribose methylation status of cap-adjacent nucleotides across animals. RNA 28:1377–1390.

Dohnalkova M, Krasnykov K, Mendel M, Li L, Panasenko O, Fleury-Olela F, Vågbø CB, Homolka D, Pillai RS. 2023. Essential roles of RNA cap-proximal ribose methylation in mammalian embryonic development and fertility. Cell Rep 42:112786.

Doitsidou M, Poole RJ, Sarin S, Bigelow H, Hobert O. 2010. *C. elegans* mutant identification with a one-step whole-genome-sequencing and SNP mapping strategy. PLoS One 5:e15435.

Dönmez G, Hartmuth K, Lührmann R. 2004. Modified nucleotides at the 5’ end of human U2 snRNA are required for spliceosomal E-complex formation. RNA 10:1925–1933.

Drazkowska K, Tomecki R, Warminski M, Baran N, Cysewski D, Depaix A, Kasprzyk R, Kowalska J, Jemielity J, Sikorski PJ. 2022. 2’-O-Methylation of the second transcribed nucleotide within the mRNA 5’’cap impacts the protein production level in a cell-specific manner and contributes to RNA immune evasion. Nucleic Acids Res 50:9051–9071.

Eijlers P, Al-Khafaji M, Soto-Martin E, Fasimoye R, Stead D, Wenzel M, Müller B, Pettitt J. 2024. A nematode-specific ribonucleoprotein complex mediates interactions between the major nematode spliced leader snRNP and its target pre-mRNAs. Nucleic Acids Res 52:7245–7260.

Ellis RE. 2022. Sex Determination in Nematode Germ Cells. Sex Dev 16:305–322.

Ewels P, Magnusson M, Lundin S, Käller M. 2016. MultiQC: summarize analysis results for multiple tools and samples in a single report. Bioinformatics 32:3047–3048.

Fasimoye RY, Spencer REB, Soto-Martin E, Eijlers P, Elmassoudi H, Brivio S, Mangana C, Sabele V, Rechtorikova R, Wenzel M, Connolly B, Pettitt J, Müller B. 2022. A novel, essential *trans*-splicing protein connects the nematode SL1 snRNP to the CBC-ARS2 complex. Nucleic Acids Res 50:7591–7607.

Feder M, Pas J, Wyrwicz LS, Bujnicki JM. 2003. Molecular phylogenetics of the RrmJ/fibrillarin superfamily of ribose 2’-O-methyltransferases. Gene 302:129– 138.

Furuichi Y, Morgan M, Shatkin AJ, Jelinek W, Salditt-Georgieff M, Darnell JE. 1975. Methylated, blocked 5 termini in HeLa cell mRNA. Proc Natl Acad Sci U S A 72:1904–1908.

Galloway A, Cowling VH. 2019. mRNA cap regulation in mammalian cell function and fate. Biochimica et Biophysica Acta (BBA) - Gene Regulatory Mechanisms 1862:270–279.

Garrison E, Marth G. 2012. Haplotype-based variant detection from short-read sequencing. arXiv:12073907 [q-bioGN].

Gene Ontology Consortium, Aleksander SA, Balhoff J, Carbon S, Cherry JM, Drabkin HJ, Ebert D, Feuermann M, Gaudet P, Harris NL, Hill DP, Lee R, Mi H, Moxon S, Mungall CJ, Muruganugan A, Mushayahama T, Sternberg PW, Thomas PD, Van Auken K, Ramsey J, Siegele DA, Chisholm RL, Fey P, Aspromonte MC, Nugnes MV, Quaglia F, Tosatto S, Giglio M, Nadendla S, Antonazzo G, Attrill H, Dos Santos G, Marygold S, Strelets V, Tabone CJ, Thurmond J, Zhou P, Ahmed SH, Asanitthong P, Luna Buitrago D, Erdol MN, Gage MC, Ali Kadhum M, Li KYC, Long M, Michalak A, Pesala A, Pritazahra A, Saverimuttu SCC, Su R, Thurlow KE, Lovering RC, Logie C, Oliferenko S, Blake J, Christie K, Corbani L, Dolan ME, Drabkin HJ, Hill DP, Ni L, Sitnikov D, Smith C, Cuzick A, Seager J, Cooper L, Elser J, Jaiswal P, Gupta P, Jaiswal P, Naithani S, Lera-Ramirez M, Rutherford K, Wood V, De Pons JL, Dwinell MR, Hayman GT, Kaldunski ML, Kwitek AE, Laulederkind SJF, Tutaj MA, Vedi M, Wang S-J, D’Eustachio P, Aimo L, Axelsen K, Bridge A, Hyka-Nouspikel N, Morgat A, Aleksander SA, Cherry JM, Engel SR, Karra K, Miyasato SR, Nash RS, Skrzypek MS, Weng S, Wong ED, Bakker E, Berardini TZ, Reiser L, Auchincloss A, Axelsen K, Argoud-Puy G, Blatter M-C, Boutet E, Breuza L, Bridge A, Casals-Casas C, Coudert E, Estreicher A, Livia Famiglietti M, Feuermann M, Gos A, Gruaz-Gumowski N, Hulo C, Hyka-Nouspikel N, Jungo F, Le Mercier P, Lieberherr D, Masson P, Morgat A, Pedruzzi I, Pourcel L, Poux S, Rivoire C, Sundaram S, Bateman A, Bowler-Barnett E, Bye-A-Jee H, Denny P, Ignatchenko A, Ishtiaq R, Lock A, Lussi Y, Magrane M, Martin MJ, Orchard S, Raposo P, Speretta E, Tyagi N, Warner K, Zaru R, Diehl AD, Lee R, Chan J, Diamantakis S, Raciti D, Zarowiecki M, Fisher M, James-Zorn C, Ponferrada V, Zorn A, Ramachandran S, Ruzicka L, Westerfield M. 2023. The Gene Ontology knowledgebase in 2023. Genetics 224. doi:10.1093/genetics/iyad031

Guo X, Zhang R, Wang J, Ding S-W, Lu R. 2013. Homologous RIG-I-like helicase proteins direct RNAi-mediated antiviral immunity in *C. elegans* by distinct mechanisms. Proc Natl Acad Sci U S A 110:16085–16090.

Haline-Vaz T, Silva TCL, Zanchin NIT. 2008. The human interferon-regulated ISG95 protein interacts with RNA polymerase II and shows methyltransferase activity. Biochem Biophys Res Commun 372:719–724.

Haussmann IU, Wu Y, Nallasivan MP, Archer N, Bodi Z, Hebenstreit D, Waddell S, Fray R, Soller M. 2022. CMTr cap-adjacent 2’-O-ribose mRNA methyltransferases are required for reward learning and mRNA localization to synapses. Nat Commun 13:1209.

Hills-Muckey K, Martinez MAQ, Stec N, Hebbar S, Saldanha J, Medwig-Kinney TN, Moore FEQ, Ivanova M, Morao A, Ward JD, Moss EG, Ercan S, Zinovyeva AY, Matus DQ, Hammell CM. 2022. An engineered, orthogonal auxin analog/AtTIR1(F79G) pairing improves both specificity and efficacy of the auxin degradation system in *Caenorhabditis elegans*. Genetics 220. doi:10.1093/genetics/iyab174

Hyde JL, Diamond MS. 2015. Innate immune restriction and antagonism of viral RNA lacking 2’-O methylation. Virology 479–480:66–74.

Inesta-Vaquera F, Chaugule VK, Galloway A, Chandler L, Rojas-Fernandez A, Weidlich S, Peggie M, Cowling VH. 2018. DHX15 regulates CMTR1- dependent gene expression and cell proliferation. Life Sci Alliance 1:e201800092.

Jiao X, Chang JH, Kilic T, Tong L, Kiledjian M. 2013. A mammalian pre-mRNA 5’end capping quality control mechanism and an unexpected link of capping to pre- mRNA processing. Mol Cell 50:104–115.

Jiao X, Doamekpor SK, Bird JG, Nickels BE, Tong L, Hart RP, Kiledjian M. 2017. 5’ End Nicotinamide Adenine Dinucleotide Cap in Human Cells Promotes RNA Decay through DXO-Mediated deNADding. Cell 168:1015–1027.e10.

Joseph BB, Blouin NA, Fay DS. 2017. Use of a Sibling Subtraction Method (SSM) for Identifying Causal Mutations in *C. elegans* by Whole-Genome Sequencing. G3 . doi:10.1534/g3.117.300135

Keith G. 1995. Mobilities of modified ribonucleotides on two-dimensional cellulose thin-layer chromatography. Biochimie 77:142–144.

Kelly WG, Xu S, Montgomery MK, Fire A. 1997. Distinct requirements for somatic and germline expression of a generally expressed *Caernorhabditis elegans* gene. Genetics 146:227–238.

Kesner JS, Chen Z, Shi P, Aparicio AO, Murphy MR, Guo Y, Trehan A, Lipponen JE, Recinos Y, Myeku N, Wu X. 2023. Noncoding translation mitigation. Nature 617:395–402.

Kiledjian M. 2018. Eukaryotic RNA 5’-End NAD+ Capping and DeNADding. Trends Cell Biol 28:454–464.

Kim D, Langmead B, Salzberg SL. 2015. HISAT: a fast spliced aligner with low memory requirements. Nat Methods 12:357–360.

Kimble J, Crittenden SL. 2007. Controls of germline stem cells, entry into meiosis, and the sperm/oocyte decision in *Caenorhabditis elegans*. Annu Rev Cell Dev Biol 23:405–433.

Krueger F. 2015. Trim Galore!: A wrapper around Cutadapt and FastQC to consistently apply adapter and quality trimming to FastQ files, with extra functionality for RRBS …. Babraham Institute.

Kruse S, Zhong S, Bodi Z, Button J, Alcocer MJC, Hayes CJ, Fray R. 2011. A novel synthesis and detection method for cap-associated adenosine modifications in mouse mRNA. Sci Rep 1:126.

Kuge H, Brownlee GG, Gershon PD, Richter JD. 1998. Cap ribose methylation of c- mos mRNA stimulates translation and oocyte maturation in Xenopus laevis. Nucleic Acids Res 26:3208–3214.

Lasda EL, Kuersten S, Blumenthal T. 2011. SL *trans*-splicing in a *Caenorhabditis elegans* in vitro extract. Cold Spring Harb Protoc 2011:db.prot5574.

Lažetić V, Batachari LE, Russell AB, Troemel ER. 2023. Similarities in the induction of the intracellular pathogen response in *Caenorhabditis elegans* and the type I interferon response in mammals. Bioessays 45:e2300097.

Li H, Durbin R. 2010. Fast and accurate short read alignment with Burrows–Wheeler transform. Bioinformatics 25:1754–1760.

Li H, Handsaker B, Wysoker A, Fennell T, Ruan J, Homer N, Marth G, Abecasis G, Durbin R, 1000 Genome Project Data Processing Subgroup. 2009. The Sequence Alignment/Map format and SAMtools. Bioinformatics 25:2078–2079.

Li Y, Lu J, Han Y, Fan X, Ding S-W. 2013. RNA interference functions as an antiviral immunity mechanism in mammals. Science 342:231–234.

Liang S, Silva JC, Suska O, Lukoszek R, Almohammed R, Cowling VH. 2022. CMTR1 is recruited to transcription start sites and promotes ribosomal protein and histone gene expression in embryonic stem cells. Nucleic Acids Res. doi:10.1093/nar/gkac122

Liao Y, Smyth GK, Shi W. 2014. featureCounts: an efficient general purpose program for assigning sequence reads to genomic features. Bioinformatics 30:923–930.

Love MI, Huber W, Anders S. 2014. Moderated estimation of fold change and dispersion for RNA-seq data with DESeq2. Genome Biol 15:550.

Mao K, Breen P, Ruvkun G. 2022. The *Caenorhabditis elegans* ARIP-4 DNA helicase couples mitochondrial surveillance to immune, detoxification, and antiviral pathways. Proc Natl Acad Sci U S A 119:e2215966119.

Mao K, Breen P, Ruvkun G. 2020. Mitochondrial dysfunction induces RNA interference in *C. elegans* through a pathway homologous to the mammalian RIG-I antiviral response. PLoS Biol 18:e3000996.

Meisel JD, Wiesenthal PP, Mootha VK, Ruvkun G. 2024. CMTR-1 RNA methyltransferase mutations activate widespread expression of a dopaminergic neuron-specific mitochondrial complex I gene. Curr Biol. doi:10.1016/j.cub.2024.04.079

Mello CC, Kramer JM, Stinchcomb D, Ambros V. 1991. Efficient gene transfer in *C. elegans*: extrachromosomal maintenance and integration of transforming sequences. EMBO J 10:3959–3970.

Mount SM, Anderson P. 2000. Expanding the definition of informational suppression. Trends Genet 16:157.

Müller MBD, Kasturi P, Jayaraj GG, Hartl FU. 2023. Mechanisms of readthrough mitigation reveal principles of GCN1-mediated translational quality control. Cell 186:3227–3244.e20.

Nance J, Frøkjær-Jensen C. 2019. The *Caenorhabditis elegans* Transgenic Toolbox. Genetics 212:959–990.

Negishi T, Kitagawa S, Horii N, Tanaka Y, Haruta N, Sugimoto A, Sawa H, Hayashi K-I, Harata M, Kanemaki MT. 2022. The auxin-inducible degron 2 (AID2) system enables controlled protein knockdown during embryogenesis and development in *Caenorhabditis elegans*. Genetics 220. doi:10.1093/genetics/iyab218

Panek J, Gang SS, Reddy KC, Luallen RJ, Fulzele A, Bennett EJ, Troemel ER. 2020. A cullin-RING ubiquitin ligase promotes thermotolerance as part of the intracellular pathogen response in *Caenorhabditis elegans*. Proc Natl Acad Sci U S A 117:7950–7960.

Perez-Riverol Y, Bai J, Bandla C, García-Seisdedos D, Hewapathirana S, Kamatchinathan S, Kundu DJ, Prakash A, Frericks-Zipper A, Eisenacher M, Walzer M, Wang S, Brazma A, Vizcaíno JA. 2022. The PRIDE database resources in 2022: a hub for mass spectrometry-based proteomics evidences. Nucleic Acids Res 50:D543–D552.

Philippe L, Pandarakalam GC, Fasimoye R, Harrison N, Connolly B, Pettitt J, Müller B. 2017. An *in vivo* genetic screen for genes involved in spliced leader *trans*- splicing indicates a crucial role for continuous *de novo* spliced leader RNP assembly. Nucleic Acids Res 45:8474–8483.

Picard-Jean F, Brand C, Tremblay-Létourneau M, Allaire A, Beaudoin MC, Boudreault S, Duval C, Rainville-Sirois J, Robert F, Pelletier J, Geiss BJ, Bisaillon M. 2018. 2’-O-methylation of the mRNA cap protects RNAs from decapping and degradation by DXO. PLoS One 13:e0193804.

Reddy KC, Dror T, Sowa JN, Panek J, Chen K, Lim ES, Wang D, Troemel ER. 2017. An intracellular pathogen response pathway promotes proteostasis in *C. elegans*. Curr Biol 27:3544–3553.e5.

Schindelin J, Arganda-Carreras I, Frise E, Kaynig V, Longair M, Pietzsch T, Preibisch S, Rueden C, Saalfeld S, Schmid B, Tinevez J-Y, White DJ, Hartenstein V, Eliceiri K, Tomancak P, Cardona A. 2012. Fiji: an open-source platform for biological-image analysis. Nat Methods 9:676–682.

Schmittgen TD, Livak KJ. 2008. Analyzing real-time PCR data by the comparative C(T) method. Nat Protoc 3:1101–1108.

Schuberth-Wagner C, Ludwig J, Bruder AK, Herzner A-M, Zillinger T, Goldeck M, Schmidt T, Schmid-Burgk JL, Kerber R, Wolter S, Stümpel J-P, Roth A, Bartok E, Drosten C, Coch C, Hornung V, Barchet W, Kümmerer BM, Hartmann G, Schlee M. 2015. A conserved histidine in the RNA sensor RIG-I controls immune tolerance to N1-2’-O-methylated self RNA. Immunity 43:41–51.

Senchuk MM, Dues DJ, Schaar CE, Johnson BK, Madaj ZB, Bowman MJ, Winn ME, Van Raamsdonk JM. 2018. Activation of DAF-16/FOXO by reactive oxygen species contributes to longevity in long-lived mitochondrial mutants in *Caenorhabditis elegans*. PLoS Genet 14:e1007268.

Shen Y, Zhang J, Calarco JA, Zhang Y. 2014. EOL-1, the homolog of the mammalian Dom3Z, regulates olfactory learning in *C. elegans*. J Neurosci 34:13364– 13370.

Simabuco FM, Pavan ICB, Pestana NF, Carvalho PC, Basei FL, Campos Granato D, Paes Leme AF, Zanchin NIT. 2019. Interactome analysis of the human Cap- specific mRNA (nucleoside-2’-O-)-methyltransferase 1 (hMTr1) protein. J Cell Biochem 120:5597–5611.

Sowa JN, Jiang H, Somasundaram L, Tecle E, Xu G, Wang D, Troemel ER. 2020. The *Caenorhabditis elegans* RIG-I homolog DRH-1 mediates the intracellular pathogen response upon viral infection. J Virol 94. doi:10.1128/jvi.01173-19

Stiernagle T. 2006. Maintenance of *C. elegans*. WormBook 1–11.

Tan X, Calderon-Villalobos LIA, Sharon M, Zheng C, Robinson CV, Estelle M, Zheng N. 2007. Mechanism of auxin perception by the TIR1 ubiquitin ligase. Nature 446:640–645.

Tecle E, Chhan CB, Franklin L, Underwood RS, Hanna-Rose W, Troemel ER. 2021. The purine nucleoside phosphorylase *pnp-1* regulates epithelial cell resistance to infection in *C. elegans*. PLoS Pathog 17:e1009350.

The UniProt Consortium, Bateman A, Martin M-J, Orchard S, Magrane M, Ahmad S, Alpi E, Bowler-Barnett EH, Britto R, Bye-A-Jee H, Cukura A, Denny P, Dogan T, Ebenezer T, Fan J, Garmiri P, da Costa Gonzales LJ, Hatton-Ellis E, Hussein A, Ignatchenko A, Insana G, Ishtiaq R, Joshi V, Jyothi D, Kandasaamy S, Lock A, Luciani A, Lugaric M, Luo J, Lussi Y, MacDougall A, Madeira F, Mahmoudy M, Mishra A, Moulang K, Nightingale A, Pundir S, Qi G, Raj S, Raposo P, Rice DL, Saidi R, Santos R, Speretta E, Stephenson J, Totoo P, Turner E, Tyagi N, Vasudev P, Warner K, Watkins X, Zaru R, Zellner H, Bridge AJ, Aimo L, Argoud-Puy G, Auchincloss AH, Axelsen KB, Bansal P, Baratin D, Batista Neto TM, Blatter M-C, Bolleman JT, Boutet E, Breuza L, Gil BC, Casals-Casas C, Echioukh KC, Coudert E, Cuche B, de Castro E, Estreicher A, Famiglietti ML, Feuermann M, Gasteiger E, Gaudet P, Gehant S, Gerritsen V, Gos A, Gruaz N, Hulo C, Hyka-Nouspikel N, Jungo F, Kerhornou A, Le Mercier P, Lieberherr D, Masson P, Morgat A, Muthukrishnan V, Paesano S, Pedruzzi I, Pilbout S, Pourcel L, Poux S, Pozzato M, Pruess M, Redaschi N, Rivoire C, Sigrist CJA, Sonesson K, Sundaram S, Wu CH, Arighi CN, Arminski L, Chen C, Chen Y, Huang H, Laiho K, McGarvey P, Natale DA, Ross K, Vinayaka CR, Wang Q, Wang Y, Zhang J. 2023. UniProt: the Universal Protein Knowledgebase in 2023. Nucleic Acids Res 51:D523–D531.

Toczydlowska-Socha D, Zielinska MM, Kurkowska M, Astha N, Almeida CF, Stefaniak F, Purta E, Bujnicki JM. 2018. Human RNA cap1 methyltransferase CMTr1 cooperates with RNA helicase DHX15 to modify RNAs with highly structured 5’ termini. Philos Trans R Soc Lond B Biol Sci 373:20180161.

Tourasse NJ, Millet JRM, Dupuy D. 2017. Quantitative RNA-seq meta-analysis of alternative exon usage in *C. elegans*. Genome Res 27:2120–2128.

Wang VY-F, Jiao X, Kiledjian M, Tong L. 2015. Structural and biochemical studies of the distinct activity profiles of Rai1 enzymes. Nucleic Acids Res 43:6596–6606.

Werner M, Purta E, Kaminska KH, Cymerman IA, Campbell DA, Mittra B, Zamudio JR, Sturm NR, Jaworski J, Bujnicki JM. 2011. 2’-O-ribose methylation of cap2 in human: function and evolution in a horizontally mobile family. Nucleic Acids Res 39:4756–4768.

Wickham H. 2016. Ggplot2: Elegant graphics for data analysis, 2nd ed, Use R! Cham, Switzerland: Springer International Publishing.

Yu G, Li F, Qin Y, Bo X, Wu Y, Wang S. 2010. GOSemSim: an R package for measuring semantic similarity among GO terms and gene products. Bioinformatics 26:976–978.

Yu G, Wang L-G, Han Y, He Q-Y. 2012. clusterProfiler: an R package for comparing biological themes among gene clusters. OMICS 16:284–287.

Yu YT, Shu MD, Steitz JA. 1998. Modifications of U2 snRNA are required for snRNP assembly and pre-mRNA splicing. EMBO J 17:5783–5795.

Zhu A, Ibrahim JG, Love MI. 2019. Heavy-tailed prior distributions for sequence count data: removing the noise and preserving large differences. Bioinformatics 35:2084–2092.

Züst R, Cervantes-Barragan L, Habjan M, Maier R, Neuman BW, Ziebuhr J, Szretter KJ, Baker SC, Barchet W, Diamond MS, Siddell SG, Ludewig B, Thiel V. 2011. Ribose 2’-O-methylation provides a molecular signature for the distinction of self and non-self mRNA dependent on the RNA sensor Mda5. Nat Immunol 12:137–143.

